# CNPY4 is a Lipid-Binding Regulator of Sphingolipid Homeostasis

**DOI:** 10.1101/2025.10.13.682155

**Authors:** Michael D. Paul, Olga Zbodakova, Isabella S. Glenn, Aleksandr Stotland, Ophir D. Klein, Natalia Jura

## Abstract

Sphingolipids function both as signaling molecules and as organizers of cell membranes, and their dysregulation has been linked to cancer, metabolic disorders, and neurodegeneration. A central node in the sphingolipid metabolism network is ceramide, which is converted into numerous derivatives, including sphingomyelin (SM). Through interactions with cholesterol, SM forms liquid-ordered microdomains that influence membrane organization and signaling. Previously, we reported that the Saposin-like (SAPLIP) protein Canopy4 (CNPY4) negatively regulates the levels of free cholesterol in the plasma membrane. Although SAPLIPs commonly regulate lipid metabolism through direct lipid interactions, CNPY4 does not bind cholesterol directly. Here, we show that CNPY4 interacts with multiple sphingolipids *in vitro*, including ceramide and SM, and with ceramide in cells. We also demonstrate that *CNPY4* knockdown elevates SM levels at the plasma membrane and disrupts cellular localization and abundance of ceramide, suggesting that the levels and consequently the homeostatic distribution of these sphingolipids is under control of CNPY4. Although the most pronounced effect of CNPY4 loss is on the ceramide/SM conversion pathway, it additionally impacts the levels of over 150 cellular lipids and modulates neutral sphingomyelinase activity, consistent with secondary disruptions in sphingolipid homeostasis. Collectively, our findings point to CNPY4 as a sphingolipid chaperone that regulates the abundance and localization of these lipids, modulating in turn cholesterol homeostasis and cellular signaling.

## Introduction

The five Canopy (*Cnpy1–5*) genes encode a family of understudied, primarily ER-localized proteins that are increasingly recognized as having important roles in health and disease^1,2^. CNPY1 has mostly been studied in zebrafish, where it plays a role in FGF signaling in the midbrain-hindbrain boundary^3^ and during Kupffer’s vesicle organogenesis^4^. The best-characterized member of the CNPY family, CNPY2, modulates the unfolded protein response by sensing ER stress, which triggers release of CNPY2 from Grp78 and subsequent binding and activation of PERK^5^. In addition, CNPY2 is important for FGF21-dependent expression of the low density lipoprotein receptor (LDLR)^6^, promotes angiogenesis^7^, and stimulates neurite outgrowth in mouse neuroblastoma cells^8^. Less is known about CNPY3, CNPY4, and CNPY5. CNPY3 (PRAT4A) and CNPY4 (PRAT4B) regulate Toll-like receptor levels at the cell surface, with CNPY3 positively regulating TLR1, whereas CNPY4 negatively regulates TLR1 but positively regulates TLR4^9–11^. CNPY4 is highly expressed in children with Kawasaki disease^12^ and skin fibroblasts from patients with Rett syndrome^13^, and its overexpression is linked to metabolic dysfunction-associated steatotic liver disease, correlating with increased risk of hepatocellular carcinoma and poor patient outcomes^14^. It has also been proposed as a biomarker of glioma progression^15^. CNPY5 (MZB1, pERp1) is important for IgM and IgA antibody secretion^16,17^, but otherwise it remains poorly characterized.

CNPY proteins are primarily composed of a saposin domain and are part of the <u>SA</u>posin-<u>L</u>ike lipid Interacting <u>P</u>rotein (SAPLIP) superfamily. CNPY3 and CNPY4 additionally contain long, highly charged C-terminal tails, predicted to be unstructured^1^. Although the functional role of the saposin domain in CNPYs remains unknown, in other SAPLIPs, these domains typically interact directly with individual lipids or membrane bilayers^18^. Through these interactions, two general mechanisms of how SAPLIPs function have been described. The first is the solubilizer model, in which a typically dimeric SAPLIP extracts a single lipid molecule from the membrane, either to present it to an enzyme or to disrupt the membrane directly through pore formation^18,19^. The second is the liftase model, in which a typically monomeric SAPLIP perturbs the membrane, allowing an enzyme’s active site to access its lipid substrate^19^. Due to these roles, SAPLIPs play important roles as lipid metabolic enzymes or their cofactors, especially in pathways involved in synthesis or metabolism of sphingolipids^18,20^.

One of the lipid classes commonly bound by saposin domains are sphingolipids, which are a diverse class of lipids with complex signaling functions. They are composed of a spingoid tail, a variable headgroup, and, except in the simplest sphingolipids such as sphingosine, an additional acyl chain tail^21^. During *de novo* sphingolipid biosynthesis, an acyl chain is added to sphingosine to form ceramide, a potent signaling and pro-apoptotic molecule^22,23^. Ceramide can then be converted into more complex sphingolipids, such as sphingomyelin (SM) and cerebrosides, depending on the modifications to its head group^21,24^. Examples of SAPLIP cofactors in sphingolipid metabolism include Saposin A (SapA), which assists galactocerebrosidase in converting galactosylceramide to ceramide^19^; Saposin D (SapD), which is a cofactor for acid ceramidase (aCDase) that converts ceramide to sphingosine^25,26^; and Saposin C (SapC), which is a cofactor for acid beta-glucosidase that converts glucosylceramide to ceramide^27^. Notably, the CNPYs closely resemble the Saposin family of SAPLIPs, with CNPY2, CNPY3, and CNPY4 being most homologous to Saposin B (SapB) and SapD, while CNPY5 is most similar to SapC.

We previously showed that *Cnpy4* knockdown in cells leads to a significant accumulation of free cholesterol at the plasma membrane, resulting in hyperactivation of Hedgehog (HH) signaling^28^. This likely reflects the central role of free cholesterol in HH pathway activation as both a post-translational modifier of the secreted signaling molecule, Sonic HH (SHH), and a direct ligand of the Smoothened receptor^29–33^. We did not detect measurable binding of cholesterol or oxysterols to the saposin domain of CNPY4^28^, suggesting that its impact on cholesterol levels is indirect.

At the plasma membrane, cholesterol exists in distinct pools: an inaccessible structural reservoir and an actively regulated exchangeable pool composed of chemically active free cholesterol and cholesterol sequestered by SM^34^. SM and cholesterol are linked from *de novo* synthesis onward, as they are co-transported from the Golgi to the plasma membrane^35^. Multiple feedback mechanisms dynamically adjust SM and cholesterol levels to maintain homeostatic balance. When SM levels in the plasma membrane decrease, cholesterol levels also decrease via increased efflux to the cytosol^36^, and the high internal cholesterol levels then inhibit cholesterol synthesis further^37^. On the other hand, excess cholesterol leads to an increase in SM synthesis^38^ and a decrease in its hydrolysis, ensuring proper cholesterol partitioning between different lipid pools at the plasma membrane^39,40^.

Given the understanding that cholesterol homeostasis is intricately interwoven with SM homeostasis, our previous findings that CNPY4 affects cholesterol in the absence of its direct binding, and the reported observations that several SAPLIP proteins regulate sphingolipid biosynthesis, we investigated the effect of CNPY4 loss on the abundance of the broader lipid landscape, with a particular focus on sphingolipids. We discovered that *Cnpy4* knockdown in cells affected global lipid homeostasis, and the pathway most affected by CNPY4 loss was the conversion of ceramide to SM. Importantly, we found that recombinant CNPY4 binds several sphingolipids, with the highest affinity measured for ceramide. Our results indicate that, by directly binding to a subset of sphingolipids, CNPY4 controls SM flux to the membrane. This indirectly impacts regulation of cholesterol membrane homeostasis, contributing to the broad effects that CNPY4 loss exerts on signaling pathways, including the HH and TLR pathways^9,10,28^.

## Results

### *Cnpy4* knockdown results in a significant shift in the cellular distribution of sphingolipids

To determine the global effects of knockdown of *Cnpy4* on cellular lipid levels, we used lipidomic mass spectroscopy (MS)^41^. Compared to a non-targeting siRNA control, *Cnpy4* knockdown over 72 hours in NIH3T3 cells, which led to maximal depletion of Cnpy4 protein (**Fig. S1**), caused significant changes in over 150 individual lipid species (out of ∼ 900 detected) across 12 different lipid classes (**Figs. 1A and Table S1**). Of these lipids, roughly 62% had decreased total levels, compared to 38% which increased. Although cholesterol was not part of this lipid panel, cholesterol esters, which are both precursors and downstream products of plasma membrane cholesterol^42,43^, were abundantly represented in the panel and sensitive to CNPY4 loss. Total cholesterol ester levels were reduced by approximately 50%, with 22 species significantly decreased upon CNPY4 loss. The decrease in cholesterol derivatives capable of conversion back to cholesterol is consistent with plasma membrane cholesterol accumulation upon *Cnpy4* knockdown, as we observed previously^28^.

**Figure 1.**
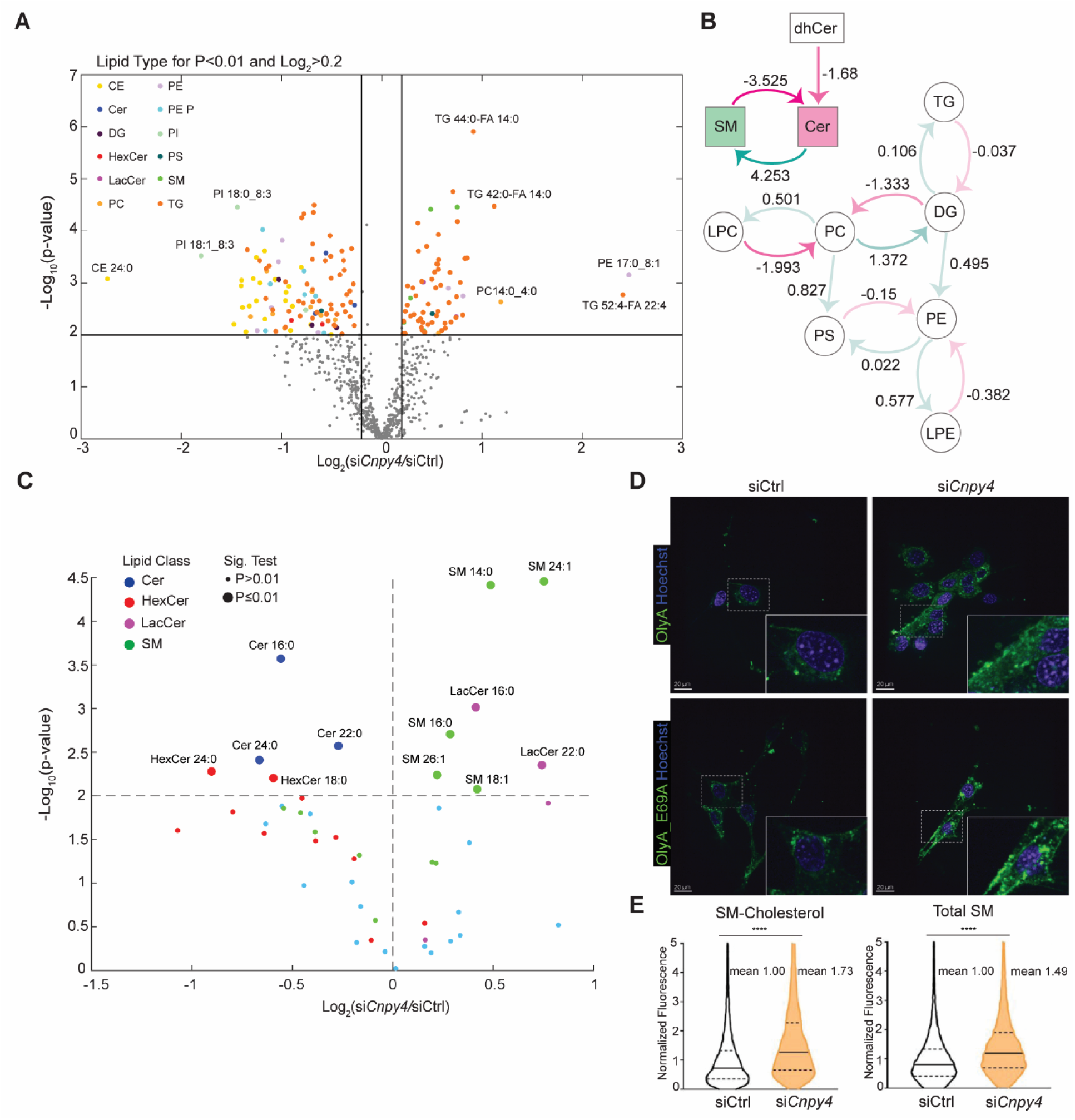
Lipidomics MS reveals broad disruption in lipid levels in the absence of CNPY4. **(A)** Volcano plot of direct infusion shotgun MS lipidomics comparing NIH3T3 cells treated with si*Cnpy4* for 72 h to non-targeting siRNA controls. Lipids with a fold change of Log_2_ > 0.2 and p < 0.01 are considered significant and shown in color, with those below either threshold shown in gray. Abbreviation and colors are as follows: CE = cholesterol ester, yellow; Cer = ceramide, blue; DG = Diacylglycerol, maroon; HexCer = hexosylceramide, red; LacCer = lactosylceramide, magenta; PC = phosphatidylcholine, goldenrod; PE = phosphatidylethanolamine (anoyl), lavender; PE P = phosphatidylethanolamine (enyl), cyan; PI = phosphatidylinositol, light green; PS = phosphatidylserine, dark green; SM = sphingomyelin, green; TG = triacylglycerol, orange. **(B)** BioPAN analysis of the lipidomics MS data. Values are Z scores: negative (magenta) indicate decreased conversion in si*Cnpy4* cells relative to control cells; positive (green) indicate increased conversion. Filled boxes mark statistical significance at p < 0.01. **(C)** Volcano plot highlighting sphingolipids from the lipidomics dataset (Cer, HexCer, LacCer, and SM), significance denotes differences between wild type and si*Cnpy4* cells at p < 0.01. **(D)** Confocal imaging of live NIH3T3 cells treated with either non-targeting siRNA control or si*Cnpy4*, and stained with either OlyA-AF488 (SM bound to cholesterol, green) or OlyA_E69A-AF488 (all SM, green) 72 h after siRNA treatment, with the nucleus stained with Hoechst (blue). The white box marks the region shown zoomed-in at lower right corner. Scale bar is 10 μm. **(E)** Violin plots depicting FACS analysis comparing NIH3T3 cells treated with either non-targeting siRNA control or si*Cnpy4*, and stained with either OlyA-AF488 or OlyA_E69A-AF488 72 h after siRNA treatment. Data are normalized to mean value of control cells. The solid line indicates the median, with the first and third quartiles indicated by the dashed lines (n = 9,410 for OlyA siCtrl, n = 9,233 for OlyA si*CNPY4*, n = 10,940 for OlyA_E69A siCtrl, and n = 13,563 for si*CNPY4* OlyA_E69A). Statistical significance was determined using a two-sided unpaired Welch’s t-test, with ****p < 0.0001. Experiments were performed at least 3 independent times.

We used BioPAN, a bioinformatic pathway analysis tool^44^, as an unbiased method to score CNPY4-dependent changes at the reaction pathway level—i.e., lipids were not considered as individual species at first, but in the context of their classes. This analysis identified interconversion between SM and ceramide as the most significantly altered pathway in the absence of CNPY4 (**Fig. 1B**). We then assessed changes at the level of individual sphingolipid species and found five SMs to be significantly increased and three ceramides to be significantly decreased upon *Cnpy4* knockdown, while the remaining SM and ceramide species did not significantly change (**Fig. 1C**). Notably, two other sphingolipid classes also exhibited uniform changes, as two hexosylceramides (HexCer) decreased and two lactosylceramide (LacCer) increased.

Individual sphingolipids have acyl chains that differ in the number of carbon atoms and saturation states. While little is known about the biological functions of individual SMs and ceramides of different chain characteristics, cells carefully control their distribution across different tissues, and their relative ratios influence the organization of plasma membrane microdomains^45–51^. We observed that *Cnpy4* knockdown altered the distribution of SM and ceramide species with varying acyl chain lengths and saturation states. The most substantial SM changes were for the 24:1, 14:0, and 16:0 chain-length variants, whose absolute mole content in the cells increased by approximately 68%, 40%, and 22%, respectively. The largest changes for ceramide were for the 24:0 and 16:0 forms, which decreased by approximately 37% and 32%, respectively. These concerted changes in sphingolipid levels across multiple species suggest that CNPY4 plays an important role in regulating metabolic flux through the sphingolipid pathway. Notably, among other significant changes observed upon *Cnpy4* knockdown, there was an increase in triacylglycerols and a decrease in phosphatidylinositol lipids (**Fig.S2**).

### Loss of CNPY4 increases the plasma membrane levels of SM

As the SM to ceramide interconversion was the most substantially impacted pathway following *Cnpy4* knockdown, we focused our analysis on this axis. Upon conversion from ceramide, SM is transported to the plasma membrane, where it becomes enriched in the outer leaflet. To investigate if the elevated global SM levels detected by lipidomics MS are associated with changed plasma membrane SM levels, we used the bacterial protein Ostreolysin A (OlyA), which specifically binds to SM. Because the wild type OlyA only recognizes SM bound to cholesterol, we also used an OlyA mutant (OlyA_E69A) that binds all SM species^52,53^. The OlyA variants, conjugated to AF488, were added to live NIH3T3 cells 72 hours after *Cnpy4* knockdown, followed by confocal imaging to assess the relative levels of both cholesterol-bound and total SM levels. We found that binding of both probes was substantially increased in cells in which *Cnpy4* was knocked down (**Fig. 1D**). To quantify this effect, we performed fluorescence-activated cell sorting (FACS) on live OlyA-labeled NIH3T3 cells, which revealed an approximately 1.5-fold increase in plasma membrane levels of cholesterol-bound and total SM following *Cnpy4* knockdown (**Fig. 1E**). Similar results were obtained in human embryonic kidney (HEK) 293 cells, indicating that the effect of CNPY4 loss on SM and cholesterol-bound SM levels is not limited to one cell type (**Fig. S3**).

### Coupled Regulation of SM and Cholesterol by CNPY4

The observed fold increase in plasma membrane SM levels is comparable to the change in plasma membrane free cholesterol upon *Cnpy4* knockdown, measured by FACS in NIH3T3 cells labeled with the bacterial derived Perfringolysin O (PFO*) probe^54^ conjugated to AF647, which we previously reported^24^. Although CNPY4 may modulate SM and cholesterol levels through distinct mechanisms, their shared metabolic and trafficking pathways suggest an interconnected mode of regulation. To assess whether such coupling occurs, we investigated the status of negative feedback loops, which are triggered by concerted SM and cholesterol trafficking to the membrane, upon *Cnpy4* knockdown.

First, we measured levels of Golgi phosphatidylinositol 4-phosphate (PI4P), which reflect the SM and cholesterol flux to the membrane through its roles in ceramide transport, SM synthesis, and cholesterol trafficking^55,56^. PI4P in the Golgi is essential for CERT function, which mediates transport of ceramide from the ER to the Golgi for conversion into SM. The conversion of ceramide to SM results in the activation of OSBP1, which then removes PI4P from the Golgi, thereby inhibiting SM synthesis, while simultaneously moving cholesterol from the ER to the Golgi, and subsequently to the plasma membrane. This means that PI4P steady-state levels are a sensitive readout of the balance between SM and cholesterol trafficking pathways and are expected to decrease in response to enhanced ceramide to SM conversion (**Fig. 2A**). This is indeed what we observed after 72 hours of *Cnpy4* knockdown by immunofluorescence imaging of NIH3T3 cells using an anti-PIP4 antibody, which revealed a ∼50% reduction in Golgi PI4P levels compared to control cells (**Figs. 2B & 2C**). This decrease in PI4P levels following CNPY4 loss was comparable to that induced by addition of exogenous ceramide, which directly stimulates SM synthesis, triggering its subsequent inhibition (**Figs. 2B & 2C**).

**Figure 2.**
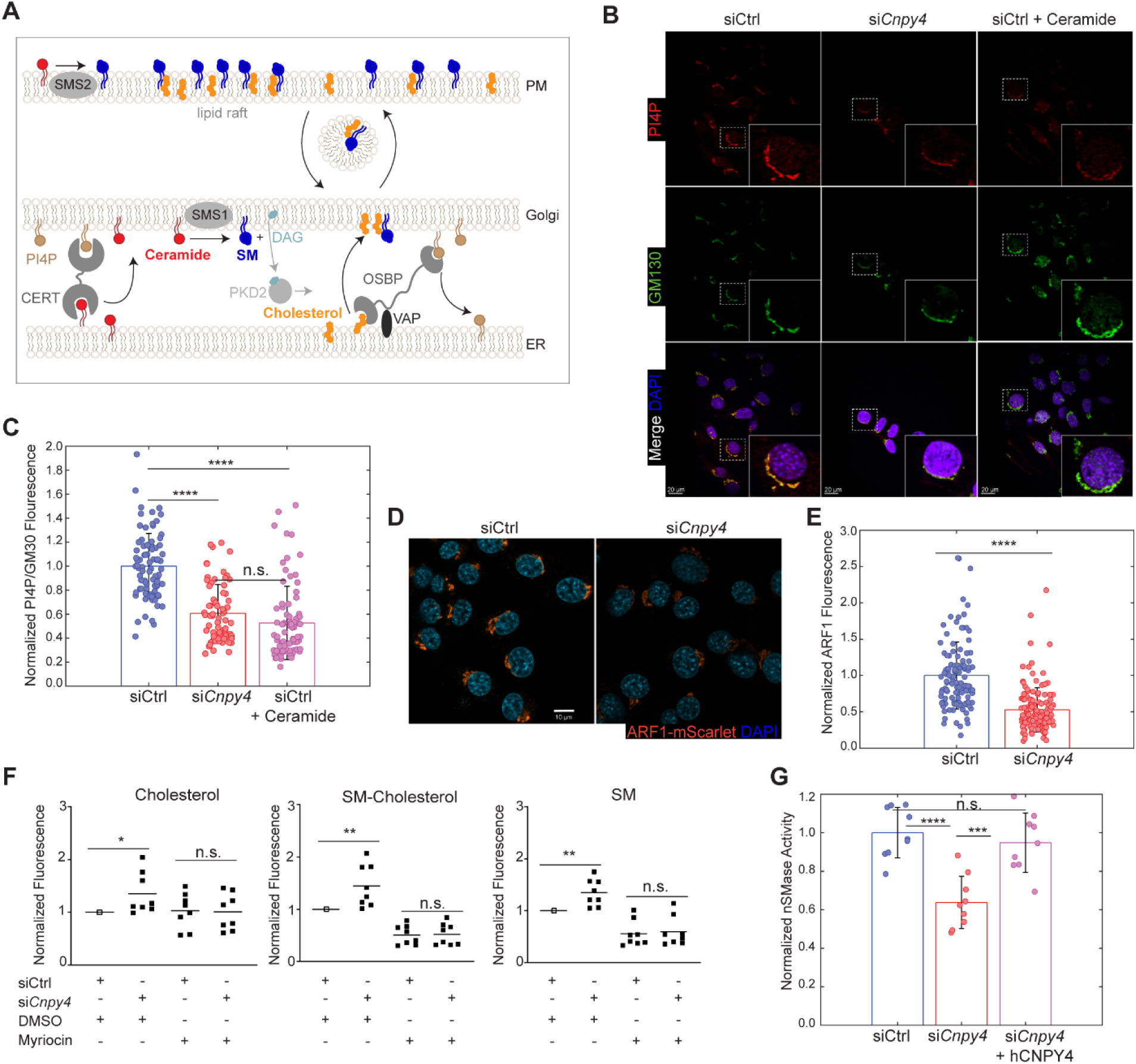
Loss of CNPY4 disrupts SM synthesis and degradation pathways. **(A)** Schematic of SM synthesis and coupled cholesterol flux. CERT transfers ER ceramide to the Golgi in a PI4P-dependent manner. SMS1 converts ceramide to SM, producing DAG, which activates PKD2, which then phosphorylates OSPB1. This OSPB1 then removes PI4P from the Golgi, and this is coupled to moving cholesterol from the ER to the Golgi, which then goes to the plasma membrane. **(B)** Confocal microscopy imaging of NIH3T3 cells treated with non-targeting siRNA control (siCtrl) alone, si*Cnpy4*, or siCtrl with 10 μM ceramide 6:0 for 30 min, fixed 72 h after siRNA treatment. PI4P signal is shown in red, GM130 in green (Golgi), and DAPI in blue (nucleus). The white box marks the region shown zoomed-in at lower right corner. Scale bar is 10 μm. **(C)** Quantification of Golgi PI4P from images in (B). PI4P/GM130 fluorescence ratios were normalized to the siCtrl mean, data are mean ± SD. Statistical significance was determined by one-way ANOVA with Tukey’s Honestly Significant Difference (n = 82 for siCtrl, n = 70 for si*Cnpy4*, and n = 81 for siCtrl + ceramide), with ****p<0.0001, n.s. p>0.5, and images were taken across 4 independent experiments. **(D)** Confocal microscopy of NIH3T3 cells with mScarlet-tagged endogenous ARF1 treated with either siCtrl or si*Cnpy4*, fixed and imaged 72 h after siRNA treatment. **(E)** Quantification of ARF1 fluorescence in cells imaged as described in (F). Fluorescence is normalized to the mean of the siCtrl cells. Error bars represent mean ± SD. Statistical significance determined by a two-sided unpaired Welch’s t-test (n = 117 for siCtrl and n = 137 for si*Cnpy4*), with ****p<0.0001, and images were taken across at least 3 independent experiments. **(F)** FACS analysis of NIH3T3 cells treated with siCtrl or si*Cnpy4* and stained for cholesterol (PFO*-AF657), SM bound to cholesterol (OlyA-AF488), and all SM (OlyA_E69A-AF488) 72 h after siRNA treatment, in the presence of 40 μM myriocin or DMSO control, where indicated. Squares represent individual experiments normalized to the mean of the corresponding siCtrl DMSO condition, with the solid bar representing the mean of all experiments. Statistical significance was determined using a one-sample t-test with a null hypothesis that the mean equals 1 (siCtrl DMSO vs si*Cnpy4* DMSO) or a two-sided unpaired Welch’s t-test (siCtrl myriocin vs si*Cnpy4* myriocin), with **p<0.01, ***p<0.001, n.s. p>0.05. (n=8 from 4 independent trials, 3,000 – 10,000 cells per point were analyzed). **(G)** Neutral sphingomyelinase activity in NIH3T3 cell lysates, 72h after transfection with either siCtrl or si*Cnpy4,* and transfected with either an empty vector control or hCNPY4. Activity was normalized to the mean of the siCtrl with empty vector control. Error bars represent mean ± SD (n = 9 from 3 independent experiments with 3 replicates per experiments). Statistical significance determined by One-way ANOVA and Tukey’s Honestly Significant Difference, with ***p<0.001, ****p<0.0001, n.s. p>0.05.

We next assessed if the decrease in PI4P levels we observed after CNPY4 loss has functional consequences on the recruitment of ADP-ribosylation factor 1 (ARF1) to the Golgi. ARF1, a G protein involved in regulation of protein and lipid trafficking^57,58^, localizes to the Golgi by binding PI4P^59^, and in turn recruits the PtdIns 4-kinase IIIβ (PI4KB/PI4KIIIβ), which generates PI4P^60–64^. We measured ARF1 levels in NIH3T3 cells endogenously expressing mScarlet labeled ARF1, and found that 72 hours after *Cnpy4* knockdown, ARF1 levels at the Golgi were substantially decreased (**Figs. 2D and 2E**). The Golgi integrity was not affected, as measured by GOLGA5 (Golgin-84) staining, indicating that the effect is specific to the PI4P pathway (**Fig. S4**). Altogether, these results are consistent with a role of CNPY4 in the control of the cholesterol/SM flux to the plasma membrane.

Since elevated SM synthesis drives cholesterol flux to the plasma membrane through the PI4P-dependent feedback loop, which we found activated upon *Cnpy4* knockdown, we hypothesized that the increase in plasma membrane cholesterol previously observed upon *Cnpy4* knockdown^28^ is driven by enhanced conversion of ceramide to SM. Based on this model, we predicted that treating cells with myriocin, a chemical inhibitor of SM biosynthesis^55,65^, should decrease membrane cholesterol levels in the absence of CNPY4. Indeed, myriocin treatment lowered free cholesterol levels at the membrane of *Cnpy4* knockdown cells to that of control cells (**Fig. 2F**). Myriocin treatment also significantly lowered levels of both cholesterol-bound and total plasma membrane SM in the *Cnpy4* knockdown cells to levels comparable to those observed in the myriocin-treated control cells (**Fig. 2F**), indicating that CNPY4 acts along the *de novo* SM synthesis pathway.

### Neutral sphingomyelinase activity is decreased after *Cnpy4* knockdown

Increased SM synthesis upon *Cnpy4* knockdown, which leads to elevated cholesterol and SM levels at the plasma membrane, and a decrease in Golgi PI4P, triggers a negative feedback loop that rapidly inhibits SM synthesis, significantly lowering SM levels within hours^55^ (**Fig. 2A**). We found that activation of the PI4P negative feedback could be measured by immunofluorescence that showed decreased Golgi PIP4 levels as quickly as 30 minutes after exogenous ceramide addition, and that it persisted 72 hours after *Cnpy4* knockdown (**Figs. 2B & 2C**). Despite the negative feedback activation, plasma membrane SM levels remained elevated 72 hours after *Cnpy4* knockdown, pointing to a potential decrease in SM degradation. Under normal conditions, SM degradation at the plasma membrane increases cholesterol ester levels as a consequence of internalization of cholesterol^42,43^. Accordingly, impaired SM degradation, induced for example by CRISPR knockout of the SM degrading enzyme neutral sphingomyelinase 2 (nSMase2)^66^, reduces cholesterol ester levels. We found that 72 hours after *Cnpy4* knockdown, cholesterol ester levels were also reduced, pointing to impaired SM degradation in the absence of Cnpy4 (**Fig. 1A**).

We therefore asked whether loss of CNPY4 resulted in decreased sphingomyelinase (SMase) activity in NIH3T3 cells. We focused on the neutral SMases (nSMases), because of their cellular localization along the ER/Golgi secretory pathway and their impact on regulation of plasma membrane SM levels^67^. We found that total nSMase activity was significantly impaired in NIH3T3 cell lysates 72 hours after *Cnpy4* knockdown compared to control cells (**Fig. 2G**). A similar effect was observed in HEK293 cells upon *CNPY4* knockdown (**Fig. S5**). Importantly, the impaired nSMase activity was rescued by exogenously expressing human CNPY4 in NIH3T3 cells following *Cnpy4* knockdown (**Fig. 2G**). To confirm that this loss of nSMase activity impaired SM degradation in live cells, we loaded NIH3T3 cells with BODIPY labeled SM and measured the fluorescent lipid levels after two hours. As expected, *Cnpy4* knockdown cells had significantly higher fluorescent signal than the control cells, acutely demonstrating the decreased rate of SM degradation (**Figs. S6A & 6B**).

Loss of nSMase activity has previously been found to increase membrane and total cholesterol levels in Jurkat cells^66^, likely because cholesterol efflux is inhibited in the absence of SM removal from the plasma membrane. To evaluate the extent of these changes relative to those induced by *Cnpy4* knockdown, we knocked down individual nSMases—*Smpd2* (nSMase1), *Smpd3* (nSMase2), or *Smpd4* (nSMase3)—in NIH3T3 cells, and measured levels of total SM and cholesterol at the cell surface. Loss of nSMase1 only had a minor effect on membrane SM levels, but loss of nSMase2 or nSMase3 resulted in a significant increase of cell surface SM (**Fig. S7A**). The increases in SM were accompanied by elevated levels of free cholesterol in the plasma membrane, to a degree similar to that observed upon *Cnpy4* knockdown (**Fig. S7B**). These findings further support the notion that CNPY4-dependent modulation of membrane cholesterol is a homeostatic response to its effect on SM anterograde flux and turnover.

### Recombinant CNPY4 binds sphingolipids but not cholesterol, *in vitro*

Several enzymes in the SM biosynthesis/degradation pathways rely on a lipid-binding saposin domain, such as acid sphingomyelinase (aSMase)^68,69^, or on a saposin domain-containing cofactor, as seen in aCDase relying on SapD^25,26^. CNPY proteins, which consist primarily of a saposin domain, do not have known lipid-binding properties, and we have previously shown that CNPY4 does not have measurable affinity for cholesterol or selected oxysterols^28^. To investigate if CNPY4 might directly bind sphingolipids, we purified recombinant full-length, N-terminally FLAG-tagged human CNPY4 (hCNPY4) (**Figs. S8A and S8B**) and tested its lipid binding properties *in vitro*, using a protein-lipid blot overlay assay containing a range of lipid classes (**Figs. S8C**). We detected measurable binding between hCNPY4 and several lipids, including ceramide 1-phosphate, phosphatidylinositol 3-phosphate (PtdIns(3)P), phosphatidic acid, and lysophosphatidic acid, but not SM. Consistent with our previous findings^28^, no binding between hCNPY4 and cholesterol was detected (**Fig. S8C**).

To assess these findings with an orthogonal approach, we used fluorescence polarization (FP) to measure binding between NBD-labeled ceramide 1-phosphate and the recombinant hCNPY4 and obtained a dissociation constant (K_d_) of 130 ± 40 nM for the interaction (**Fig. S8D**). Using FP, we also measured the binding between hCNPY4 and ceramide, which was not included in the protein-lipid blot overlay assay, and obtained a comparably high binding affinity, with a K_d_ of 80 ± 25 nM (**Fig. 3A**). Interestingly, although hCNPY4 did not bind SM displayed on the lipid array, it bound SM with detectable, albeit weaker affinity in the FP assay (280 ± 120 nM) (**Fig. 3B**). No binding could be detected between hCNPY4 and cholesterol (**Fig. S8E**).

**Figure 3.**
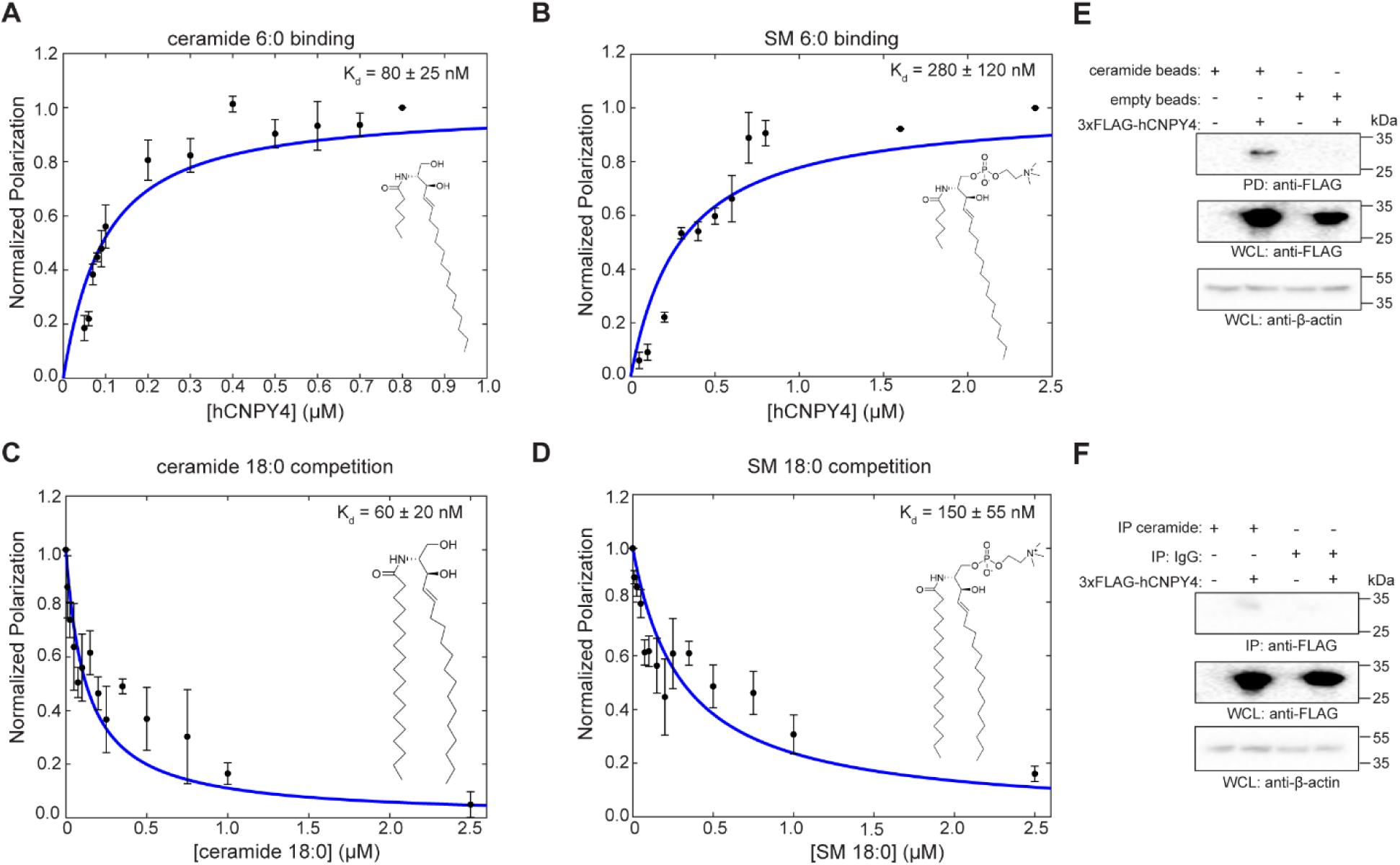
CNPY4 binds ceramide *in vitro* and in cells. **(A-D)** Fluorescence polarization (FP) binding curves of recombinant N-terminally FLAG-tagged hCNPY4 to the indicated lipids. (A-B) FP was measured over increasing hCNPY4 concentration in the presence of (A) 25 nM NBD labeled ceramide 6:0, and (B) 25 nM NBD labeled SM 6:0. (C) FP competition assay with 25 nM NBD labeled ceramide 6:0, 150 nM hCNPY4, and increasing concentrations of unlabeled ceramide 18:0. (D) FP competition assay with 25 nM NBD labeled SM 6:0, 300 nM hCNPY4, and increasing concentrations of unlabeled SM 18:0. Curves were normalized with the baseline (no protein) set to 0 and the highest concentration tested set to 1. Curve fitting and K_d_ determination are described in Methods. Data points represent the mean ± SEM from at least 3 independent experiments, with technical triplicate averaged per experiment considered one replicate. Inserted images represent respective lipid molecules (fluorophore not shown). **(E)** Pulldown (PD) of transiently transfected FLAG–hCNPY4 from NIH3T3 whole cell lysates (WCLs) using streptavidin coated beads conjugated to biotinylated ceramide 6:0. Cells transfected with either an empty vector (EV) or FLAG–hCNPY4 and a PD with unconjugated streptavidin coated beads were used as negative controls. **(F)** Immunoprecipitation (IP) of transiently transfected FLAG–hCNPY4 from NIH3T3 from whole cell lysates with an anti-ceramide antibody. Cells transfected with an empty vector (EV) or FLAG–hCNPY4 and an IP with an anti-IgG were used as negative controls. Representative Western blots from 3 independent experiments are shown in (E-F).

In the FP experiments, we used commercially available sphingolipids bearing short, saturated six carbon (6:0) acyl chains with an NBD fluorophore conjugated to the acyl chain. Because these lipids are considerably shorter than the long-chain species typically present in cells^70,71^, we performed competition FP binding assays using the 18:0 forms of ceramide and SM, which are among the most abundant species present in mammalian membranes^70,71^, to compete with the labeled probes. The ceramide 18:0 displaced the NBD-labeled ceramide 6:0 with an apparent K_d_ of 60 ± 20 nM (**Fig. 3C**), while SM 18:0 displaced the NBD-labeled SM 6:0 with an apparent K_d_ of 125 ± 45 nM (**Fig. 3D**).

### CNPY4 binds ceramide in cells

To investigate whether CNPY4 interacts with ceramide in a cellular context, we used biotinylated ceramide (6:0) conjugated to streptavidin-coated beads to pulldown CNPY4 directly from cell lysates. The FLAG-tagged human CNPY4 transiently expressed in NIH3T3 cells was robustly precipitated by the immobilized ceramide, while no protein was detected when unconjugated streptavidin-coated beads were used as a control (**Fig. 3E**). We also tested the interaction between CNPY4 and endogenous ceramides, using an anti-ceramide antibody. Exogenously expressed hCNPY4 in NIH3T3 cells clearly coimmunoprecipitated with ceramide, while no interaction was seen with an IgG pulldown control (**Fig. 3F**). Since the anti-ceramide antibody is specific for ceramide and the headgroup is the only unique portion of ceramide, these results suggest that CNPY4 binds to the acyl and/or sphingosine tail of ceramide, to allow formation of the ternary complex. This mode of lipid binding is consistent with how other SAPLIPS, particularly members of the Saposin family, recognize their lipid ligands^18,19^. It also offers a mechanistic explanation for the broader specificity of CNPY4 to sphingolipids, as these lipids share a common tail architecture but have unique head groups.

### Loss of CNPY4 transiently increases Golgi ceramide levels

Our findings suggest a model in which CNPY4 acts as a chaperone for ceramide, restricting its availability for conversion to SM. This model is supported by two key observations: first, CNPY4 binds ceramide with high affinity *in vitro* and in cells; second, after *Cnpy4* knockdown, SM and cholesterol levels are increased at the plasma membrane, and the PI4P-depleting feedback loop is activated—both hallmarks of increased ceramide to SM conversion. In this model, acute loss of CNPY4 would be expected to transiently raise ceramide levels, which would later decline as ceramide is converted to SM and cholesterol accumulates. We therefore examined cellular ceramide levels at early and late time points following *Cnpy4* knockdown.

We used immunofluorescence to assess ceramide levels in NIH3T3 cells over time after *Cnpy4* knockdown (**Figs. 4A & 4B**). In support of our hypothesis, ceramide levels increased significantly in the Golgi as early as 16 hours after *Cnpy4* knockdown. At this time, no significant increase in plasma membrane levels of SM could be measured (**Figs. S9**). SM levels gradually increased at the plasma membrane over time and reached a maximum 72 hours post knockdown, at which point Golgi ceramide was decreased, suggesting its conversion to SM. Further evidence for the disruption of ceramide localization in cells in the absence of CNPY4 comes from the analysis of the distribution of 6:0 ceramide fluorescently labeled with NBD exogenously added to NIH3T3 cells (**Fig. 4C**). Thirty minutes after ceramide addition, *Cnpy4* knockdown cells had the same total amount of NBD-ceramide as control cells, but there was increased Golgi accumulation, supporting the model in which CNPY4 binding ceramide regulates its flux through an anabolic pathway towards SM production (**Fig. 4D & 4E**). We hypothesize that this increase in ceramide-to-SM conversion, observed at later timepoints after *Cnpy4* knockdown, might represent a compensatory mechanism to remove the excessive ceramide to counteract cellular toxicity that stem from its accumulation.

**Figure 4.**
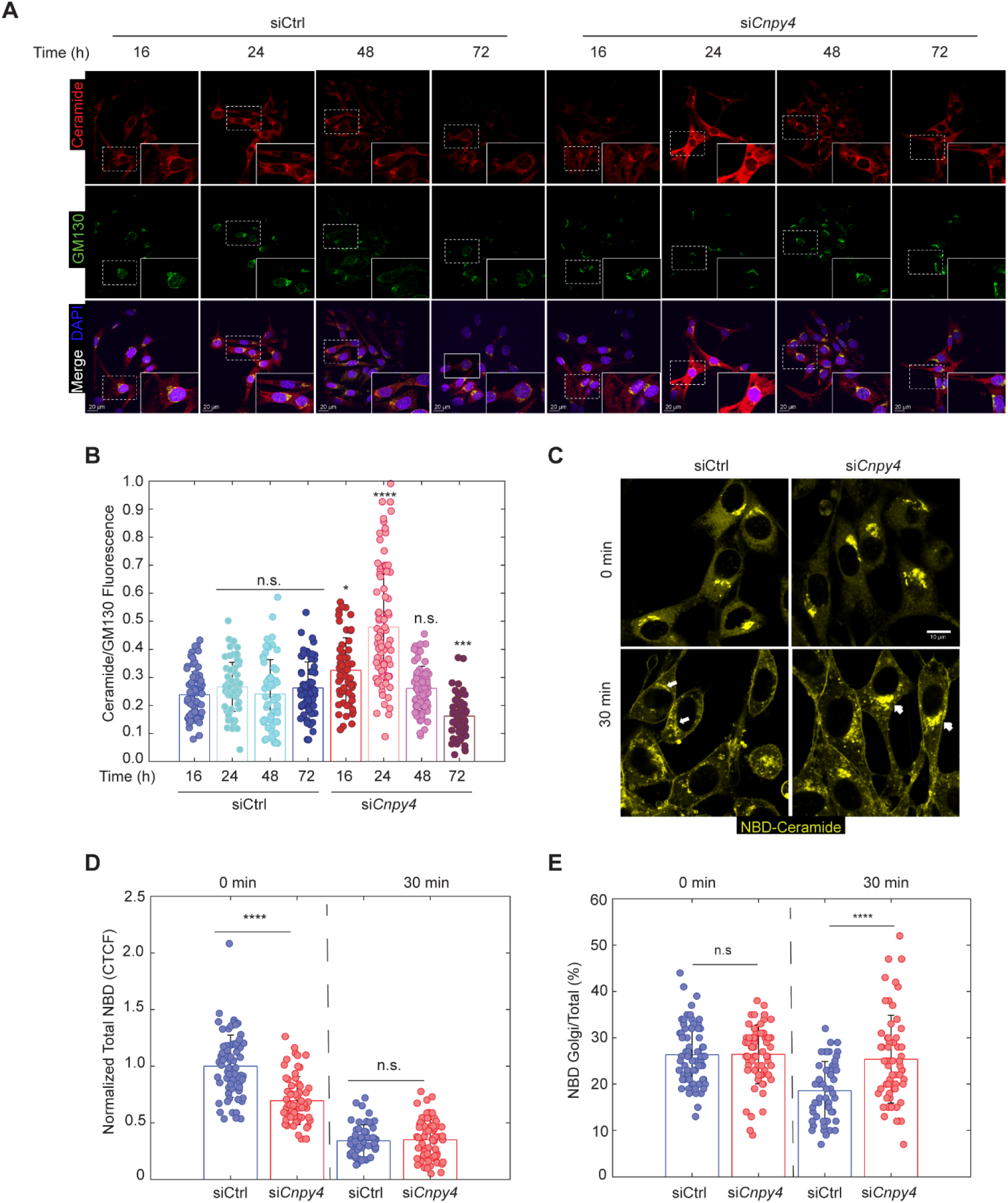
*Cnpy4* knockdown disrupts ceramide levels and localization. **(A)** Confocal imaging of NIH3T3 cells treated with either non-targeting siRNA (siCtrl) control or si*Cnpy4*, followed by fixation and staining for ceramide (red) with an anti-ceramide antibody, Golgi (green) with an anti-GM130 antibody, and the nucleus (blue) with DAPI. The white box marks the region shown zoomed-in at lower right corner. Scale bar is 10 μm. **(B)** Quantification of Golgi ceramide in cells imaged as described in (A). Ratio of ceramide to GM130 fluorescence is shown (mean ± SD). Statistical significance was determined by One-way ANOVA and Tukey’s Honestly Significant Difference (n = 74 for siCtrl 16 h, n = 72 for siCtrl 24 h, n = 63 for siCtrl 48 h, n = 70 for siCtrl 72 h, n = 65 for si*Cnpy4* 16h, n = 84 for si*Cnpy4* 24h, n = 77 for si*Cnpy4* 48h, and n = 62 for si*Cnpy4* 72h), and shown relative to siCtrl 16 h condition, with *p<0.05, *P<0.001, ****p<0.0001, n.s. p>0.5. Images were taken across 3 independent experiments. **(C)** Confocal microscopy of NIH3T3 cells treated with either siCtrl or si*Cnpy4*, incubated with 1 μg/mL fluorescently tagged NBD-ceramide 6:0 for the indicated time intervals, and imaged 72 h after siRNA treatment. The white arrows point to ceramide accumulated in the Golgi. Scale bar is 10 μm. **(D)** Quantification of total NBD fluorescence in cells imaged as described in (C). Fluorescence was normalized to the mean of the time 0 in siCtrl-treated cells. Error bars represent mean ± SD. Statistical significance was determined by a two-sided unpaired Welch’s t-test (n=55 siCtrl 0 min, n=55 si*Cnpy4* 0 min, n=60 siCtrl 30 min, and n=52 si*Cnpy4* 30 min), with ****p<0.0001, n.s. p>0.05. **(E)** Quantification of the percentage of total NBD fluorescence localized to the Golgi in the same cells as in (D). Error bars represent mean ± SD. Statistical significance was determined by a two-sided unpaired Welch’s t-test (n = 55 for siCtrl 0 min, n = 55 for si*Cnpy4* 0 min, n = 60 fro siCtrl 30 min, and n = 52 for si*Cnpy4* 30 min), with ****p<0.0001, n.s. p>0.05.

### Gene expression changes in the absence of CNPY4 are consistent with decreased ceramide levels, although ceramide synthesis enzymes are largely unaffected

Tight regulation of ceramide levels is critical for maintaining cellular homeostasis. The changes in ceramide levels over time following *Cnpy4* knockdown suggest broader cellular rewiring may occur to compensate for ceramide dysregulation. To address this, we performed RNA-seq to measure global gene expression changes in NIH3T3 cells 72 hours after *Cnpy4* knockdown.

Analysis of genes involved in ceramide metabolism (**Fig. S10A**) showed changes consistent with decreased ceramide levels. Expression of *Asah1*, which encodes aCDase that degrades ceramide specifically at the lysosome, was significantly increased (**Fig. S10B**). Genes encoding other lysosomal sphingolipid degrading enzymes were also increased, including *Gba* (lysosomal acid glucosylceramidase), *Smpd1* (aSMase/sphingomyelin phosphodiesterase), and *Sphk1* (sphingosine kinase 1), which degrade HexCer, SM, and sphingosine, respectively. Upregulation of lysosomal enzymes that degrade sphingolipids would reduce overall SM and other sphingolipid levels and restore lipid homeostasis, while avoiding additional ceramide production in the Golgi or ER, where its generation could be problematic in the absence of CNPY4. We also observed reduced levels of *Ugcg* (ceramide glucosyltransferase), the enzyme that converts ceramide into HexCer, consistent with the decrease in HexCers measured by lipidomics MS (**Fig. 1D**). Further changes in ceramide metabolic enzymes were seen in the significant increase of *Cerk* (ceramide kinase), which converts ceramide to ceramide 1-phosphate, and the decrease of *Sms2* (sphingomyelin synthase 2), which converts ceramide to SM at the plasma membrane (**Fig. S10C**). On the other hand, *Sms1*, which converts ceramide to SM at the Golgi, was unaffected. Genes involved in *de novo* synthesis of ceramide in the ER, such as those encoding the ceramide synthases (*Cers1-5*) and sphingolipid delta(4)-desaturase DES1 (*Degs1*), were also unaffected (**Fig. S10D**). Taken together, the gene expression changes support a role for CNPY4 as an important homeostatic regulator of ceramide levels, with its loss leading to a rewiring of ceramide metabolic pathways. More broadly, Gene Ontology (GO) analysis identified that one of the areas with the largest number of differentially expressed genes was control of lipid localization, with an increase in expression of several cholesterol transporter genes. Interestingly, genes encoding for regulators of extracellular matrix components, calcium and metal ion homeostasis, and axon development were also significantly altered (**Fig.S10E**).

## Discussion

In most mammalian cell types, SM composes 10-20 mol% of the plasma membrane, and through its interactions with cholesterol, it plays a central role in plasma membrane domain organization, contributing to the formation of liquid-ordered microdomains, also known as lipid rafts^34,72–74^. Here, we demonstrate that CNPY4 plays a key role in maintaining SM levels at the plasma membrane, and link these findings to our previous observation that *Cnpy4* knockdown increases the levels of total cellular cholesterol and free cholesterol at the plasma membrane^28^.

We propose that these effects are part of the same mechanism, as the synthesis, degradation, and trafficking of SM and cholesterol are tightly coupled, with perturbations in one lipid typically mirrored in the other^35–40^. Consequently, *Cnpy4* knockdown has no effect on cholesterol levels when SM synthesis is inhibited. Also consistent with coupling between SM and cholesterol, *Cnpy4* knockdown activates the PI4P-OSBP1 feedback that links ER-to-PM cholesterol transport to SM synthesis^55,56^.

We also found that *Cnpy4* knockdown changes the distribution of individual SMs in cells, as well as other sphingolipids including ceramide. Cells contain sphingolipids with many different chain lengths, and different cells and tissues have different distributions for a given sphingolipid^70,71^. Although the physiological importance of these differences in chain length is still poorly understood, they have different biophysical properties, distinct effects on the organization of the plasma membrane, and differing specificity for lipid-binding proteins^45–51^. For example, cells engineered to only have SM 24:0 had a dramatically reduced presence of lipid rafts compared to cells only containing SM 16:0^51^. Broadly reducing levels of SMs with a chain length greater than 20 caused a decrease in both the plasma membrane lipid order and the cholesterol levels^46^. The CNPY4-dependent SM chain-length changes are thus unlikely to be incidental, and testing their functional consequences is an important next step.

Although CNPY4 and other CNPY family members have a saposin domain, which is known to bind a range of lipids in other SAPLIP proteins, CNPYs have not previously been shown to interact with lipids. We provide evidence for high affinity binding to ceramide (K_d_ = 80 ± 25 nM) and SM (K_d_ = 155 ± 55 nM). These affinities match those of known sphingolipid binders, including CERT (K_d_ = 58 nM for ceramide 6:0)^75^, and cytosolic phospholipase A2 (K_d_ = 15–125 nM for ceramide-1-phosphate)^76–78^. Collectively, these observations suggest that CNPY4 acts as a sphingolipid chaperone and point to a potentially closer connection between CNPY proteins and Saposin proteins, to which CNPYs are most closely similar in primary sequence. Saposins are known to chaperone many sphingolipids, including SapA binding to galactosylceramides^19^, SapB binding to cerebroside and SM^79^, SapC^27^ binding to glucosylceramides, and SapD binding to ceramide^25,26^.

Together, our findings are consistent with a model in which CNPY4 regulates the conversion of ceramide to SM at the ER/Golgi interface (**Fig. 5**). CNPY4 has a noncanonical, weaker ER-retention sequence (REDL instead of KDEL)^1^, and is hence detected across the ER/Golgi compartment^1,28^. We observed that when CNPY4 is depleted, ceramide—both endogenous and a fluorescently labeled version added exogenously—transiently accumulates in the Golgi, potentially due to impaired localization or impaired presentation to the enzymes that convert it into downstream sphingolipids. As accumulation of ceramide results in cell toxicity^80–83^, one way for cells to adapt is to lower ceramide levels by increasing SM synthesis, resulting in long term decreased levels of ceramide in the Golgi and increased SM at the plasma membrane. In parallel, reduced nSMase activity diminishes the back-conversion of SM to ceramide and, because SM degradation normally accompanies internalization of plasma-membrane cholesterol^36^, this helps sustain the rise in surface cholesterol. Increased SM flux to the plasma membrane is also coupled to enhanced ER to plasma membrane cholesterol transport via the PI4P-OSBP1 feedback loop^35,55,56^. We see all these homeostatic mechanisms engaged upon CNPY4 loss. Furthermore, by seventy-two hours, the cell reinforces this rebalancing through a transcriptional response aimed at restoring lipid homeostasis. We observed upregulation of mRNA encoding for proteins that would lower ceramide levels by converting it to other sphingolipids, such as *Cerk*, and lysosomal sphingolipid-degrading enzymes, such as *Spmd1* and *Asah1*, suggesting a possible route by which cells reduce SM levels at the plasma membrane without generating ceramide at the ER or Golgi. Additionally, we found elevated mRNAs encoding for enzymes involved in cholesterol transport, and significant changes in transcripts for genes associated with lipid localization in general, such as the ATP-binding cassette transporters *Abca9* and *Abca8b*, the oxysterol binding protein *Osbp2*, and the high-density lipoprotein component *ApoD*. Together, these data underscore the impact CNPY4 has on cellular lipid homeostasis.

**Figure 5.**
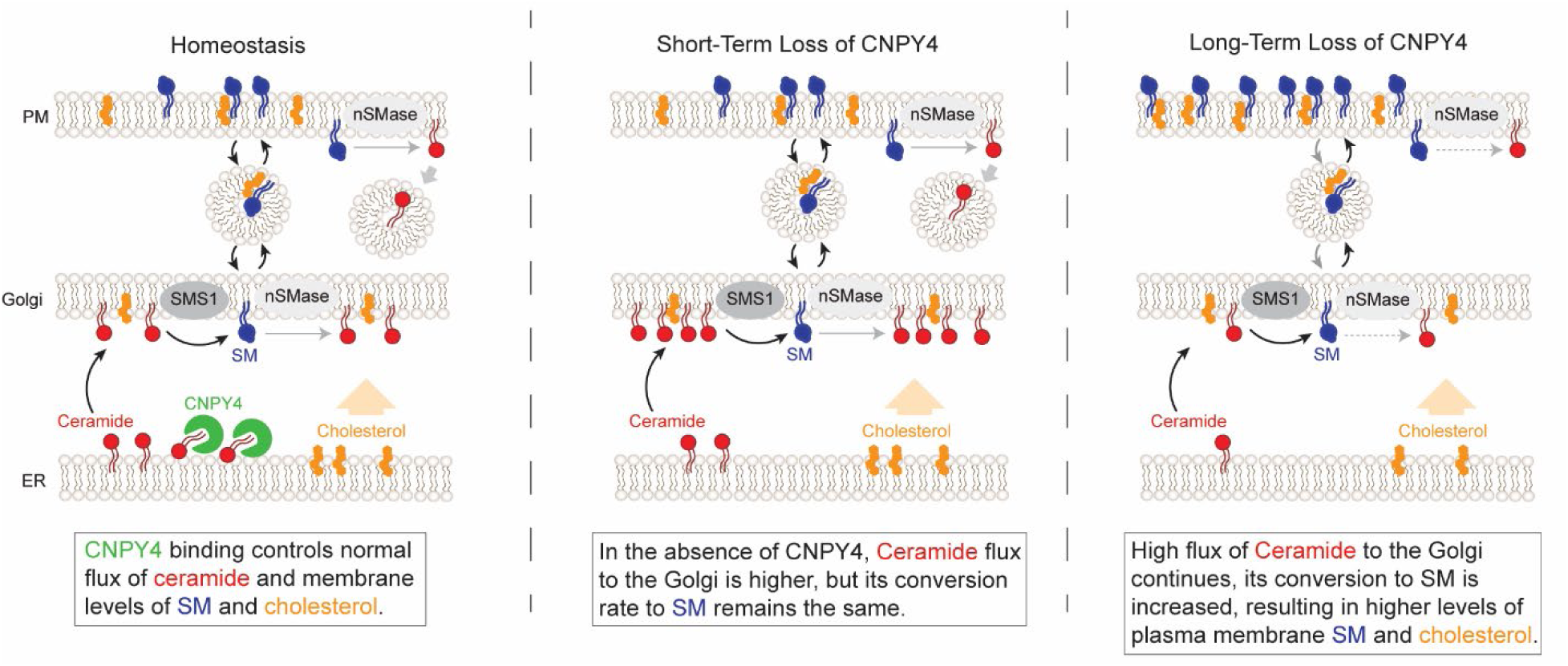
Proposed model for the role of CNPY4 in regulation of ceramide and SM. During homeostasis, CNPY4 binds to ceramide, thereby controlling its flux to the Golgi and conversion to SM (left panel). Shortly after the loss of CNPY4, free ceramide is now transported to Golgi more efficiently, but its conversion to SM is not increased, leading to a temporary ceramide accumulation in the Golgi (middle panel). Prolonged loss of CNPY4 leads to increased conversion of ceramide to SM, which is then transported to the plasma membrane. This results in the accumulation of plasma membrane cholesterol, keeping the membrane SM:cholesterol ratio constant, and decreased ceramide in the Golgi. We also observe decreased nSMase activity upon CNPY4 loss, additionally contributing to the increase in SM levels (right panel).

Although CNPY4 binds several lipids *in vitro*, its cellular specificity for ceramide is likely shaped by its subcellular localization to the ER/Golgi network. Given the diversity of lipids in the ER and Golgi, many of which are more abundant than ceramide, CNPY4 is likely to have additional lipid ligands, whose identities and *in vivo* binding conditions will require further investigation. CNPY4’s broad lipid engagement may stem from the binding mode itself, though in the absence of direct structure studies, we can only speculate. The anti-ceramide antibody which we used to immunoprecipitate CNPY4 from cells is predicted to bind to the unique sphingolipid headgroup, which implies that CNPY4’s lipid recognition must center on the aliphatic tails; otherwise, it would not be able to engage in a ternary complex with the ceramide-bound antibody. Notably, this is the same binding mode as seen in other SAPLIPs, which is unsurprising given their roles as cofactors that present sphingolipids to enzymes modifying the lipid headgroup. An example is seen in the SapA dimer, which forms a tetrameric complex with two β-galactocerebrosidase monomers, where SapA binds the glycosphingolipid tail and positions its headgroup for cleavage at the enzyme’s active site^19^. A similar lipid tail-bind mechanism is seen in the crystal structure of SapB bound to a fluorescently labeled globotriaosylceramide and of SapB in complex with α-galactosidase A^84^. A distinct example is surfactant protein B, a phospholipid binding and transfer SAPLIP essential for the for the formation of lamellar bodies, which by binding to phospholipid tails promotes lateral interactions between acyl chains^85^.

By influencing membrane SM and cholesterol membrane content, especially SM bound to cholesterol, CNPY4 may influence the organization of liquid-ordered microdomains, providing a unifying model for its diverse effects on signaling. Pathways reported to depend on CNPY4, including SHH signaling^28^, Toll-like receptor signaling^9,10^, and fibroblast growth factor (FGF) receptor signaling^28^, all rely on cholesterol-and SM-rich microdomains for proper function. In SHH signaling, free cholesterol both activates the Smoothened receptor and post-translationally modifies the SHH ligand^29–33^. Toll-like receptor membrane abundance and signaling decrease when SM synthesis is inhibited^86^, as these receptors are enriched in liquid-ordered microdomains^87,88^. FGF2 secretion is dependent on cholesterol and SM^89,90^, and FGF receptor 2 and its cofactors are enriched in liquid-ordered microdomains^91,92^. Thus, the pleiotropic effects of CNPY4 on these pathways likely arise from its ability to modulate membrane lipid composition and, in turn, the formation of signaling microdomains.

Dysregulated liquid-ordered microdomains are linked to defects in immunity^93–95^, cancer progression^96–98^, cardiovascular disease^99,100^, and neurodegeneration^101^, aligning with emerging roles of CNPY4 in human diseases^2^. *CNPY4* mRNA levels are elevated in children with Kawasaki disease^12^ and sepsis^102^, and cellular stress increases CNPY4 protein levels in macrophages^103^. Consistently, sphingolipid homeostasis is essential for immune system function, and its disruption contributes to autoimmune disease^23,104,105^. Beyond the immune system, higher *CNPY4* levels correlate with worse glioma outcomes^15^, whereas lower levels of CNPY4 are seen in Alzheimer’s disease patients^106^. The brain’s unique SM and ceramide distribution^70,71^, together with sphingolipids/nSMase dysregulation in neurodegeneration^23,67^, positions CNPY4 as a potential modulator of central nervous system pathology. Defining CNPY proteins’ molecular mechanisms of regulating sphingolipid homeostasis will be key to uncovering their signaling impact under normal conditions and in disease, and will inform therapeutic targeting.

## Methods

### Expression vectors and cell culture

Human CNPY4 constructs containing a 3xFLAG tag directly after the signal sequence were synthesized by Genscript, subcloned into a pcDNA3.1 vector (Thermo Fisher Scientific), and verified by DNA sequencing (Elim Biotechnology). NIH3T3 and HEK293 cells (ATCC) were cultured in high glucose Dulbecco’s Modified Eagle Medium (DMEM) (Thermo Fisher Scientific) supplemented with 10% fetal bovine serum (FBS) (Cytiva) and 1% penicillin-streptomycin (Cytiva) and incubated at 37 °C with 5% CO_2_. NIH3T3 ARF1-mScarlet reporter cells were generated by CRISPR/Cas9-mediated homology-directed repair. An mScarlet fluorescent tag was inserted in-frame at the C-terminus of the endogenous *Arf1* locus, following the last exon, using a donor construct containing a 3xGGGS linker and homology arms of approximately 200 – 300 bp. Correctly targeted clones were isolated by FACS sorting and verified by sequencing. Routine mycoplasma testing was performed on all cell lines using the MycoAlert mycoplasma detection kit (Lonza). Effects of exogenous lipid exposure were minimized by culturing cells in reduced-serum OPTI-MEM (Thermo Fisher Scientific) prior to experiments (24 h for NIH3T3 cells, 4 h for HEK293 cells). To inhibit SM synthesis, cells were seeded in medium supplemented with 40 μM myriocin (Cayman Chemical Company) 24 h before siRNA transfection, and the myriocin-containing media was changed daily until completion of the experiment 72 h post transfection.

### Transfection

NIH3T3 and HEK293 cells were seeded 24 h prior to being transiently transfected at ∼50 – 60% confluency using 22.5 pmol of gene-specific siRNA SMARTpool or siGENOME Non-Targeting siRNA Pool #1 (Dharmacon) using lipofectamine RNAiMax (Thermo Fisher Scientific) according to the manufacturer’s protocol for 16 – 72 h before analysis. hCNPY4 was transiently overexpressed in NIH3T3 cells at ∼70 – 80% confluency 48 h before cell lysis using lipofectamine LTX with Plus reagent (Thermo Fisher Scientific) according to the manufacturer’s protocol. Efficacy of both knockdown and overexpression were assessed by Western blotting.

### qRT-PCR analysis and bulk RNA sequencing

For qRT-PCR, total cellular RNA was isolated by direct lysis of NIH3T3 cells on tissue culture plate using GenCatch™ Total RNA Mini Prep Kit (Epoch Live Science). Reverse transcription was done using High-Capacity cDNA Reverse Transcription Kit (Applied Biosystems) according to the manufacturer’s protocol, using 500 ng of RNA per reaction. Primers were designed using Primer-BLAST software^107^ and primer efficiency was verified to be within the range of 90 – 110%. qPCR reaction was conducted in 10 μl volume using 2x MasterMix (Bio-Rad) on Bio-Rad CFX384 cycler. C_t_s were normalized to the mean of *Gapdh* and *Rpl19*. To minimize DNA contamination, total RNA for bulk RNA sequencing was isolated using NucleoSpin RNA Plus XS (Macherey-Nagel). Sequencing, data normalization, and GO analysis was done by Novogene.

### Immunofluorescence

NIH3T3 cells were seeded at 1.0 x 10^5^ cells per dish into 35 mm collagen coated glass bottom petri dishes (MatTek), one day prior to a specific cell treatment. *Cnpy4* knockdown was performed as described in the “siRNA and plasmid transfection” subsection. For ceramide treatment, cells were supplemented with 10 μM of ceramide 6:0 (Milipore Sigma, 376650) or equivalent volume of DMSO (0.1% of total volume) 30 min prior to cell fixation. Cells were rinsed twice with PBS and then fixed with 3.7% formaldehyde for 10 min at room temperature with gentle rocking. Immediately following fixation, cells were rinsed three times with PBS and incubated with the blocking/permeabilization buffer (PBS, 2.5% BSA, 0.1% Triton X-100) for 20 min at room temperature with gentle rocking. Cells were then stained with a primary antibody (anti-GM130, 1:750 [rabbit, Thermo Scientific, PA5-95727 or MA5-35107] and either anti-PtdIns(4)P [anti-PI4P], 1:500 [mouse, Echelon Biosciences, Z-P004] or anti-Ceramide, 1:500 [mouse, Enzo Life Sciences, ALX-804-196-T050]), diluted in the blocking/permeabilization buffer, for 1 h at room temperature with gentle rocking. Next, cells were washed three times with PBS and stained with a secondary antibody (AlexaFluor 488 anti-rabbit IgG, 1:1000 [donkey, Life Technologies, A21206] or AlexaFluor 647 anti-mouse IgG [goat, Life Technologies, A21236]) for 1 h at room temperature with gentle rocking. Prolong Gold AntiFade with DAPI (Cell Signaling Technologies) was applied to stain the nucleus before sealing with a glass cover slip. Images were acquired an Eclipse Ti with a CSU-X1 spinning disc confocal (Nikon) and Clara interline CCD camera (Andor) with a Plan Apo 60x oil objective (Nikon) running Nikon Elements 5.02 build 1266 (Nikon).

### Live cell fluorescent lipid experiments

Two days following siRNA transfection, NIH3T3 cells were seeded onto glass-bottom plates (Mattek) to reach approximately 50% of confluency for imaging the following day. On the subsequent day, cells were incubated with either 5 μg/mL of BODIPY™ FL C12-SM (Thermo Fisher Scientific) or 1 μg/ml NBD ceramide 6:0 (Avanti Polar Lipids). The incubation periods were 45 min for BODIPY™ FL C12-SM and 30 min for NBD ceramide 6:0. Ceramide staining was conducted at room temperature to prevent ceramide metabolization. After staining, the medium was replaced with OPTI-MEM, and cells were incubated at 37 °C for the specified time duration and then cooled to 4 °C to stop metabolic reactions. Imaging was performed at room temperature using a Zeiss LSM900 confocal microscope equipped with an Airyscan2 detector. A Plan Apo LD 40x objective with water immersion was used for imaging. For each frame, 10 Z-sections, spanning a total of 9 μm, were captured.

### Image analysis

Immunofluorescence-stained and live-cell fluorescence images were analyzed using FIJI v1.53f51. For immunofluorescence, the Golgi was manually outlined based on the GM130 staining (AF488 channel), and then the fluorescence intensity of GM130, PI4P (AF647 channel), or ceramide (AF647 channel) was measured in the region of interest. The post background corrected ratio of PI4P or ceramide to GM130 was then calculated for each cell. Analogously, in experiments tracking NBD-ceramide, the Golgi was manually outlined based on ceramide accumulation proximal to nucleus and cell boundaries were outlined based on the fluorescence signal of ceramide (NBD channel) or SM (BODIPY channel). NBD or BODIPY fluorescence was measured in regions of interest as a sum across the Z-stack. Each condition was imaged in at least three independent experiments, and the data from all cells across all experiments were combined. Statistical significance was determined using either a two-sided unpaired Welch’s t-test or a One-way ANOVA followed by Multcompare using Tukey’s Honestly Significant Difference in MATLAB 2023b (MathWorks).

### OlyA protein expression and purification

The plasmids encoding for the 6xHis tagged OlyA proteins were obtained from Dr. A. Radhakrishnan (University of Texas Southwestern Medical Center) and were purified as described previously^53,108^. In brief, the plasmids were transformed in BL21 (DE3) pLysS *E. coli* competent cells, and starter culture was grown overnight at 37 °C while shaking at 220 rpm. 10 mL of starter culture was used to inoculate 1L of culture, which was grown at 37 °C while shaking at 220 rpm, before being induced with 1 mM IPTG, and then further grown at 18 °C for 16 h. Cells were pelleted by centrifugation with a JA 8.5 rotor (Avanti) at 4,000 x g at 4 °C for 40 min. Pellets were flash frozen for later purification or resuspended in binding buffer (50 mM Tris-HCl pH 7.5, 150 mM NaCl, 1 mM TCEP, and 1% SDS) supplemented with 1 mg/mL lysozyme, 0.4 mg/mL PMSF, and a complete EDTA-free protease inhibitor cocktail minitablet (Roche). Cells were lysed using sonication and clarified using an Avanti centrifuge equipped with a JLA 25.50 rotor at 25,000 x *g* for 1 hour at 4 °C. Clarified lysates were loaded into a Ni-NTA resin column and the column was washed with 10 column volumes of binding buffer supplemented with 50 mM imidazole followed by elution of bound proteins with a 50 – 300 mM imidazole gradient. Protein-rich purified fractions were concentrated using an Amicon Ultra-15 3k MWCO centrifugal filter (Millipore Sigma) and labeled with Alexa Fluor 488 C5-maleimide dye (Thermo Fisher Scientific) according to the manufacturer’s protocol. Free dye was removed by buffer exchange into SDS-free binding buffer supplemented with 300 mM imidazole using an Amicon Ultra-15 3k MWCO centrifugal filter (Millipore Sigma). The His-tag was cleaved by incubating with lab-purified TEV protease (10:1 OlyA:TEV) for 16 h at 4 °C with gentle rocking. Cleaved OlyA was separated from uncleaved protein using Ni-NTA column, and the flowthrough was concentrated with an Amicon Ultra-15 3k MWCO centrifugal filter (Millipore Sigma), and further purified by size exclusion chromatography using a Superdex 200 Increase 16/600 GL column (GE Life Sciences) in an SDS-free binding buffer. The protein was concentrated with an Amicon Ultra-15 3k MWCO centrifugal filter (Millipore Sigma), and flash frozen in liquid nitrogen, followed by storage at -80 °C until used.

### PFO* and OlyA staining and FACS analysis

NIH3T3 cells were harvested 16 h, 24 h, 48 h, and 72 h post-siRNA transfection. Cells were washed with FACS buffer (HBSS, 1% BSA, 10 mM HEPES) and stained in parallel with 5 μg/ml PFO*-AF647, OlyA-AF488, or OlyA_E69A-AF488 for 30 min on ice in the dark. Following staining, cells were rinsed and resuspended in FACS buffer supplemented with DAPI. Fluorescence was measured using a LSRFortessa flow cytometer (BD Biosciences) or an SH800 Cell Sorter (Sony). Data analysis was performed using FlowJo^TM^ v10 software (BD Life Sciences). Mean fluorescence was calculated from cells gated for singlets (FSC-A vs. FSC-H) and live cells (DAPI-negative) and background fluorescence of unstained cells was subtracted.

### Neutral sphingomyelinase activity assay

Neutral sphingomyelinase activity of cell lysates was determined using the Colorimetric Sphingomyelinase Assay Kit (Millipore Sigma) according to the manufacture’s protocol. In brief, NIH3T3 cells were seeded in six-well plates at 1.5 x 10^5^ cells per well 24 h prior to siRNA knockdown and treated with the indicated condition. Cells were washed twice with ice-cold PBS prior to addition of lysis buffer (150 mM NaCl, 50 mM Tris pH 8.0, 0.5% Triton X-100, 0.5% NP-40, and cOmplete mini EDTA-free protease inhibitor cocktail [Roche]) for 45 min on ice. For each condition, 50 μL of sample was added to a clear, flat-bottom 96-well plate along with 50 μL of the sphingomyelin working solution from the kit, and then incubated at 37 °C for 1 h. Next, 50 μL of the sphingomyelinase assay mixture from the kit was added to each sample, followed by incubation for at least 2 h at room temperature in the dark. Absorbance was measured at 655 nm. Total protein was quantified by BCA assay and equalized across conditions (Millipore Sigma). Each condition was measured at least three independent times, and statistical significance was determined using One-way ANOVA followed by Multcompare using Tukey’s Honestly Significant Difference in MATLAB 2023b (MathWorks).

### hCNPY4 protein expression and purification

Expi293F suspension cells (Thermo Fisher Scientific) were maintained at 37 °C and 8% CO_2_ while rocking at 125 rpm in 125 mL non-baffled sterile cap vented flasks. The N-terminally 3xFLAG-tagged CNPY4 (hCNPY4) construct was transfected into 30 mL of Expi293F culture at a cell density of 4.0 x 10^6^ cells/mL using ExpiFectamine 293 transfection reagent (Thermo Fisher Scientific) according to the manufacturer’s protocol at 1 μg/mL. Enhancer was added 18 h post-transfection, and cells were collected 48 h post-enhancer addition in an Allegra X-14 centrifuge (Beckman Coulter) at 1,000 x g for 10 min. Pellets were flash frozen in liquid nitrogen for later purification or resuspended in binding buffer (50 mM Tris-HCl pH 7.5, 150 mM NaCl, 2 mM MgCl_2_) with 1 mM NaF, 1 mM Na_3_VO_4_, DNase I, 1 mM PMSF, 4 mM benzamidine, a complete EDTA-free protease inhibitor cocktail minitablet (Roche), and a Pierce protease inhibitor minitablet (Thermo Fisher Scientific) supplemented fresh. If frozen, pellets were thawed on ice and homogenized by gentle pipetting. Cells were lysed using an EmulsiFlex-C5 homogenizer (Avestin) at 10,000 – 15,000 psi, and the cell lysate was spun down in an Avanti centrifuge with a JA 25.50 rotor at 20,000 x g for 40 min, at 4 °C. The clarified lysate was then incubated with 125 μL of G1 Flag resin (Genscript) per 30 mL culture overnight at 4 °C with rotating before washing with 50 bead volumes of lysis buffer. The resin was then incubated with 10 bead volumes of elution buffer (lysis buffer + 0.25 mg/mL FLAG peptide [SinoBiological]) for 1 h at 4 °C with rotating and protein was eluted by gravity flow. The eluate was concentrated using an Amicon Ultra-0.5 10k MWCO centrifugal filter (Millipore Sigma), before being loaded and separated on a Superdex 200 Increase 10/300 GL column (GE Life Sciences). hCNPY4 peak fractions were collected, concentrated, and flash frozen.

### Lipid strips

10 ng of purified hCNPY4 was spotted onto a Mega Lipid Strip (Echelon Biosciences) as a positive control. The strip was then blocked in blocking buffer (TBS-T with 3% BSA) for 1 h at room temperature with gentle rocking. Next the strip was incubated with 8 μg/mL of hCNPY4 in blocking buffer for 1 h at room temperature with gentle rocking. After washing three times with TBS-T for 5 min each with gentle rocking at room temperature, the strip was incubated with primary antibody (anit-CNPY4, 1:500 [rabbit, Novus Biologicals, NBP1-81085] at room temperature with gentle rocking. Following an additional three washes with TBS-T for 5 min each with gentle rocking at room temperature, the strip was incubated with secondary antibody (anti-rabbit, 1:10000 [goat, Cell Signaling, 7074S]). ECL prime (VWR) were used for detection.

### Fluorescence polarization

Purified hCNPY4 in buffer (50 mM Tris-HCl pH 7.5, 150 mM NaCl) was analyzed for binding to 25 nM NBD-ceramide-1-phosphate 6:0 (Echelon Biosciences, S-500N6), 25 nM NBD-ceramide 6:0 (Avanti Polar Lipids, 810209P), 25 nM NBD-sphingomyelin 6:0 (Avanti Polar Lipids, 810218P), and 50 nM BODIPY-cholesterol (Cayman Chemical, 24618) at the indicated protein concentrations. Experiments were performed with a 20 μL reaction volume in triplicate using a flat, black-bottom 384-well plate (Corning) on a BioTek Synergy H1 plate reader using a kinetic read of fluorescent polarization with filters (ex 485/20 nm, Em 528/20 nm, top 510 nm) running Gen5 3.10 (BioTek). The reaction was incubated for 10 min at room temperature prior to measurements. Data fitting was done using MATLAB 2023b (MathWorks) to equation (1)^109^

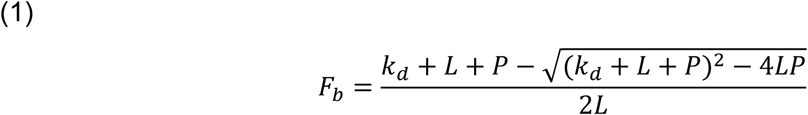

where F_b_ is the fraction of labeled lipid bound to protein (i.e., the normalized fluorescent polarization measurement), K_d_ is the protein-lipid dissociation constant, L is the total concentration of fluorescently labeled lipid, and P is the total concentration of protein.

Binding of hCNPY4 to ceramide 18:0 (Avanti Polar Lipids, 860518P) and SM 18:0 (Avanti Polar Lipids, 860586P) was tested using a competition assay against the corresponding 6:0 NBD-lipid. Experiments were performed in the same way as with the labeled NBD-lipid binding above, except the hCNPY4 concentration was fixed (150 μM for testing ceramide 18:0 and 300 μM for testing SM 18:0), the unlabeled lipid was at the indicated concentration, and the reaction was incubated for 20 min at room temperature prior to measuring the absorbance. The hCNPY4 concentration was selected to be low enough to avoid the plateau region which would be insensitive to the competitor, and high enough to provide good signal as the labeled lipid is displaced^110^. Data fitting was done using MATLAB 2023b (MathWorks) to equation (2)^111^.

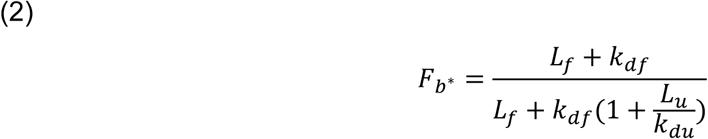

where F_b*_ is the fraction of labeled lipid bound to protein relative to the maximal amount which can be bound given the concentration of protein present (i.e., the normalized fluorescent polarization measurement), K_df_ is the protein-lipid dissociation constant for the fluorescently labeled lipid (measured in a separate experiment described above), L_f_ is the total concentration of the fluorescently labeled lipid, K_du_ is the protein-lipid dissociation constant for the unlabeled lipid, and L_u_ is the protein-lipid dissociation constant for the unlabeled lipid. For all fluorescence polarization experiments, hCNPY4 binding to each lipid was tested at least three independent times using hCNPY4 from two separate protein purifications.

### Co-immunoprecipitation and ceramide pulldown

NIH3T3 cells were seeded in a 150 mm petri dish at 5.0 x 10^6^ cells per dish and transfected the following day. Cells were washed twice with ice-cold PBS prior to addition of lysis buffer (150 mM NaCl, 50 mM Tris pH 8.0, 0.5% Triton X-100, 0.5% NP-40, 1 mM NaF, 1 mM Na_3_VO_4_, 1 mM EDTA, and cOmplete mini EDTA-free protease inhibitor cocktail). To ensure complete lysis, cells were scraped and incubated with rotation at 4 °C for 45 min. Lysates were clarified by centrifugation for 10 min at 21,000 x g at 4 °C. The clarified lysates were pre-cleared with either Protein A beads (Invitrogen, 101042) (for Co-IP) or streptavidin magnetic beads (New England Biolabs, S1420S) (for ceramide pulldown) for 30 min at 4 °C with rotation. Protein A beads were conjugated to 10 μg of anti-ceramide antibody (mouse, Enzo Life Sciences, ALX-804-196-T050), and the complex was incubated with the cell lysate at 4 °C with rotation overnight. Streptavidin magnetic beads were conjugated to 6:0 biotin-ceramide (Echelon Biosciences, S-300B) such that the ceramide concentration was 10 μM, and the complex was incubated with the cell lysate at 4 °C with rotation overnight. The beads were washed three times with lysis buffer, and then the bound protein was eluted by boiling in SDS-loading buffer for 10 min, followed by analysis by Western blot.

### Lipidomics mass spectroscopy

NIH3T3 were seeded in 150 mm plates at 3.5 x 10^6^ cells per plate and transfected the following day with either a non-targeting siRNA control (siCtrl) or si*Cnpy4* 72 h prior to harvesting. Cells were detached with trypsin, pelleted, resuspended in DMEM and counted to ensure 5 x 10^6^ cells per pellet, resuspend and rinsed three times with PBS, and then pelleted and flash frozen in liquid nitrogen. Cell pellets from 4 separately transfected replicates were generated per condition. Lipids were extracted from cell pellets using the BUME method^112^. Briefly, cell pellets were suspended in 500 μL of 3:1 v/v butanol/methanol and homogenized with the Retsch MixerMIll 301 at 25Hz for 5 min. 500 μL of 3:1 v/v heptane/ethyl acetate and 500 μL of 1% acetic acid in water was added to the tube and the lysate was homogenized again at 25Hz for 5 min. Samples were then spun for 10 min at 4000 x g and 600 μl of upper phase was transferred to a new Eppendorf tube. 500 μL of heptane/ethyl acetate was added to the lower phase and homogenization was repeated as above. The samples were centrifuged, and the 600 μL of upper phase was combined with the previously transferred supernatant. Internal standards for lipidomics were added to the extract, and sample was dried under nitrogen gas, and resuspended in 250 μL of 10 mM ammonium acetate in 1:1 v/v dichloromethane:methanol.

Lipids were profiled using The Sciex Lipidyzer platform^113^. The system consists of a QTRAP 5500 with SelexION Ion Mobility Technology and a Shimadzu Nexera X2 system consisting of a controller (CMB-20A), two HPLC pumps (LC-30AD), autosampler (SIL-30AC), and column oven (CTO-20A). The pumps were configured to run an isocratic running solution). The autosampler was configured for a 50 μL infusion. The system was controlled by Analyst software (version 1.6.3). Shotgun Lipidomics Assistant software v 1.5^114^ was used to analyze and quantify lipid species. Lipid concentrations were determined by the SLA software using the ratio of the endogenous lipid to internal standard.

### LION and BioPAN analysis

Lipid ontology (LION) analysis was performed in ranking mode using the default settings with the online server (http://www.lipidontology.com)^115,116^. Bioinformatics Methodology For Pathway Analysis (BioPAN) was performed with the online server (https://www.lipidmaps.org/biopan) with the LipidLynxX^117^ option to convert the lipid names to a compatible nomenclature and a statistical significance threshold of 0.01^44,118^.

## Data and code availability

This paper does not report original code. Any additional information required to reanalyze the data reported in this paper is available from the lead contact upon request.

## Acknowledgments

We thank M. Kinnebrew and R. Rohatgi for the PFO* probe, E. Linossi for assistance with purifying the OlyA probes, H. Torosyan for assistance with purifying hCNPY4, and P. Marangoni for technical support. We thank the members of the Jura and Klein labs for helpful discussions. Confocal imaging was performed in part at the UCSF Center for Advanced Light Microscopy, which is supported by the UCSF Research Evaluation and Allocation Committee, the Gross Fund, and the Heart Anonymous Fund. We thank the Cedars-Sinai Proteomics and Metabolomics Core facility for supporting the lipidomics data collection and analysis. This work was supported by UCSF CVRI T32 HL 7731-29 training grant to M.D.P, R01-DE033576 to O.D.K., and the Program for Breakthrough Biomedical Research at UCSF to N.J. and O.D.K.

## Author Contributions

Experiments were conceptualized by M.D.P, O.Z., O.D.K, and N.J. Experiments were performed by M.D.P. and O.Z. I.S.G. assisted with purification of hCNPY4 and the *in vitro* lipid binding assays. A.S. performed the lipidomics MS experiments and assisted with the analysis of the data. M.D.P., O.Z., O.D.K., and N.J. wrote the manuscript, with edits from I.S.G. and A.S. The project was supervised by O.D.K and N.J.

