## Supplemental Figures for "CNPY4 is a Lipid-Binding Regulator of Sphingolipid Homeostasis"

| Abbreviation | Full Lipid Class Name | # Changed, p<0.01 (Increase, Decrease) |
|---|---|---|
| CE | Cholesterol ester | 22 (0,22) |
| Cer | Ceramide | 3 (0,3) |
| DG | Diacylglycerol | 3 (0, 3) |
| FFA | Free fatty acid | n/a |
| HexCer | Hexosylceramide | 2 (0,2) |
| LPC | Lysophosphatidylcholine | 0 |
| LPE | Lysophosphatidylethanolamine | 0 |
| LacCer | Lactosylceramide | 2 (2,0) |
| PA | Phosphatidic acid | n/a |
| PC | Phosphatidylcholine | 10 (2,8) |
| PE | Phosphatidylethanolamine (anoyl) | 8 (3,5) |
| PE P | Phosphatidylethanolamine (enyl) | 7(0,7) |
| PG | Phosphatidylglycerol | 0 |
| PI | Phosphatidylinositol | 3 (0,3) |
| PS | Phosphatidylserine | 2 (1,1) |
| SM | Sphingomyelin | 5 (5, 0) |
| TG | Triacylglycerol | 107 (53, 54) |

**Table S1 Summary of the lipidomics MS analysis.** The first column lists the abbreviation for each class of lipid and the second the full lipid class name. The third column indicates the total number of individual lipids within the lipid class whose concentration were significantly ($p<0.01$) different in si*Cnpy4* cells versus siCtrl cells. Numbers in parentheses are the number of indivdual lipid species which changed (first increased, second decreased) in the absence of CNPY4. Note that the FFAs and Pas were excluded from analysis, as they were detected in the blanks.

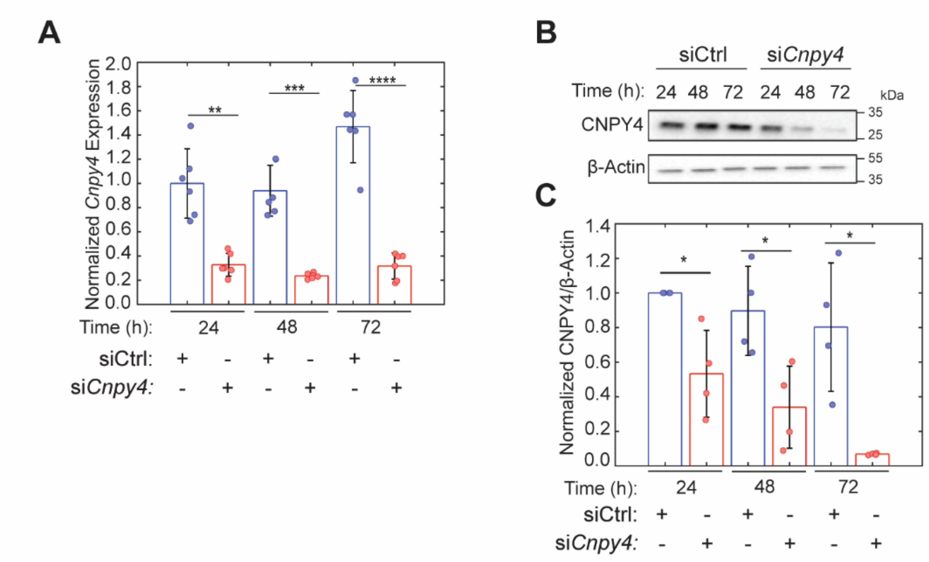


**Figure S1 Efficiency of *Cnpy4* knockdown over time. (A)** qPCR analysis of *Cnpy4* mRNA levels in NIH3T3 cells treated with either a non-targeting siRNA control (siCtrl) or si*Cnpy4* for the indicated time. Error bars represent the mean ± SD (n = 6, from 3 independent trials with two replicates per experiment). Statistical significance was determined for each time point comparing siCtrl to the si*Cnpy4* conditions using a two-sided unpaired Welch's t-test, with **p<0.01, ***p<0.001, ****p<0.0001 **(B)** Representative Western blots showing levels of CNPY4 and β-actin (loading control) in NIH3T3 cells treated with either siCtrl or si*Cnpy4* for the indicated time. **(C)** CNPY4 abundance normalized to β-actin obtained by densitometry measurements of Western blot signals. Each experiment was normalized to the ratio for the 24 h siCtrl condition. Error bars represent the mean ± SD from 4 independent experiments. Statistical significance was determined for each time point comparing siCtrl to the si*Cnpy4* conditions, using a one-sample t-test with a null hypothesis that the mean equals 1 (24 h) or two-sided unpaired Welch's t-test (48 h and 72 h), with *p<0.05.

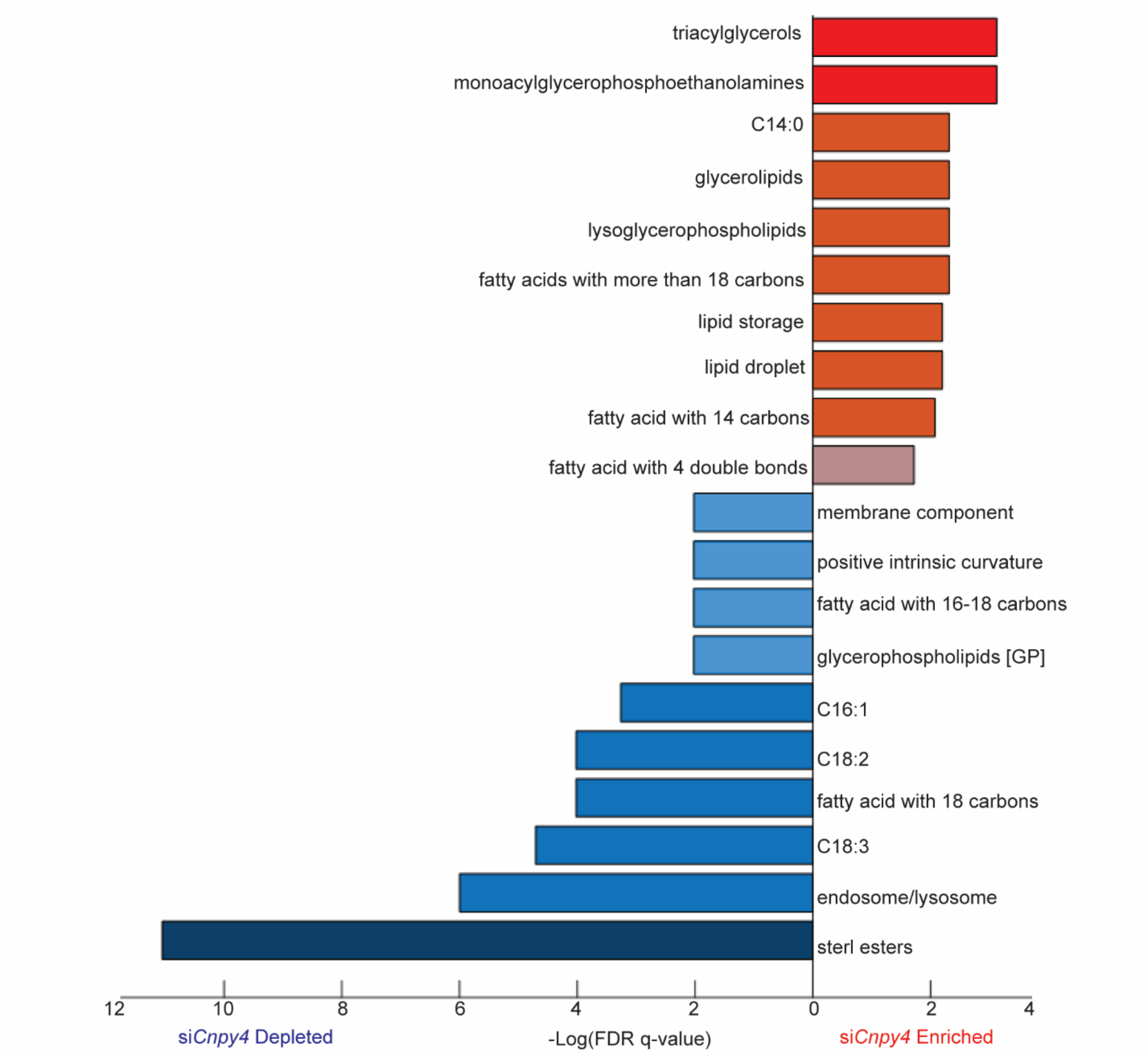


**Figure S2 LION analysis of lipidomics MS.** Most statistically significant changes from the LION analysis are shown. In LION analysis, individual lipids are annotated as having specific properties belonging to four major categories: “lipid classification,” “chemical and physical properties,” “function,” and “sub-cellular component.” Individual lipids are annotated with as many properties as apply to that lipid. Lipid properties that decreased in the absence of CNPY4 are shown in blue, and those that increased are shown in red.

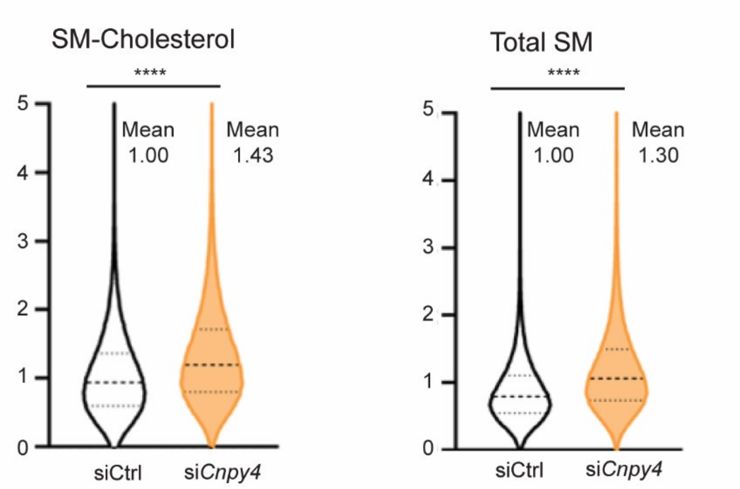


**Figure S3 Effect of *CNPY4* knockdown on plasma membrane SM levels in HEK293 cells.** Violin plots representing FACS analysis of HEK293 cells treated with either non-targeting siRNA control (siCtrl) or si*CNPY4*, and stained with either OlyA-AF488 (SM bound to cholesterol, left) or OlyA_E69A-AF488 (all SM, right) 72 h after siRNA treatment. Data were normalized to the mean signal of the control cells. The solid line indicates the median, with the first and third quartiles indicated by the dashed lines (n = 226,033 for OlyA siCtrl, n = 219,239 for OlyA si*CNPY4*, n = 245,307 for OlyA_E69A siCtrl, and n = 249,156 for si*CNPY4* OlyA_E69A). Statistical significance was determined using a two-sided unpaired Welch's t-test, with ****$p<0.0001$. Experiments were performed 3 independent times with similar results.

A

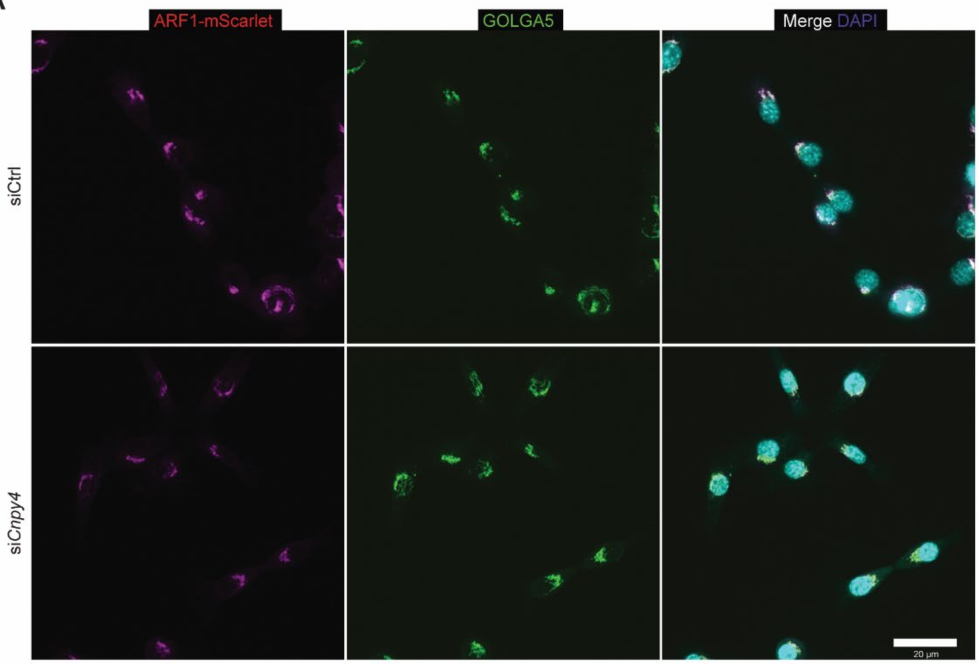


B

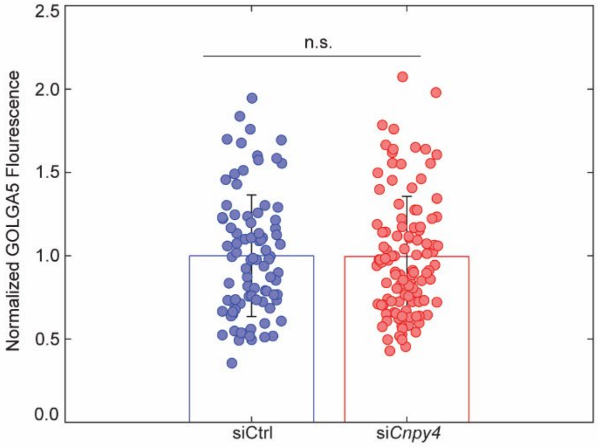


**Figure S4 GOLGA5 levels upon *Cnpy4* knockdown in NIH3T3 cells. (A)** Confocal microscopy of NIH3T3 cells with mScarlet-tagged endogenous ARF1 treated with either siCtrl or si*Cnpy4*, fixed 72 h after siRNA treatment, stained with an anti-GOLGA5 antibody (green), and imaged. **(B)** Quantification of the background-corrected total cell fluorescence in the green channel (i.e., the GOLGA5 fluorescence) in cells imaged as described in (A). Statistical significance was determined with a two-sided unpaired Welch's t-test (n=86 for siCtrl and n=107 for si*Cnpy4*), with n.s. $p>0.05$.

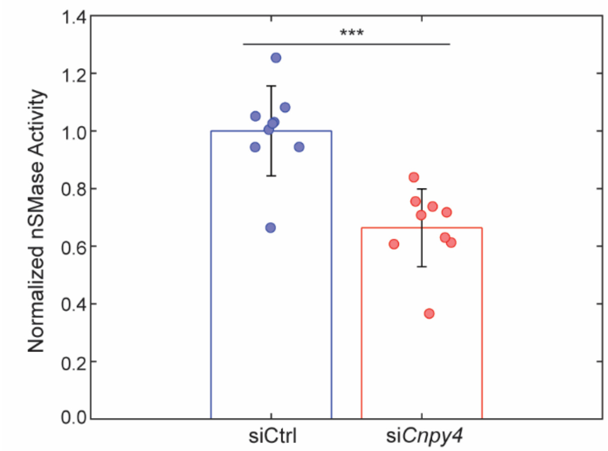


**Figure S5 nSMase activity in HEK293 cells upon *CNPY4* knockdown.** nSMase activity measured in HEK293 cell lysates 72 h post treatment with either non-targeting siRNA control (siCtrl) or si*CNPY4*. Activity was normalized to the mean of the siCtrl condition. Error bars represent mean ± SD (n = 9 from 3 independent experiments with three replicates per experiment). Statistical significance was determined using a two-sided unpaired Welch's t-test, with ***p<0.001.

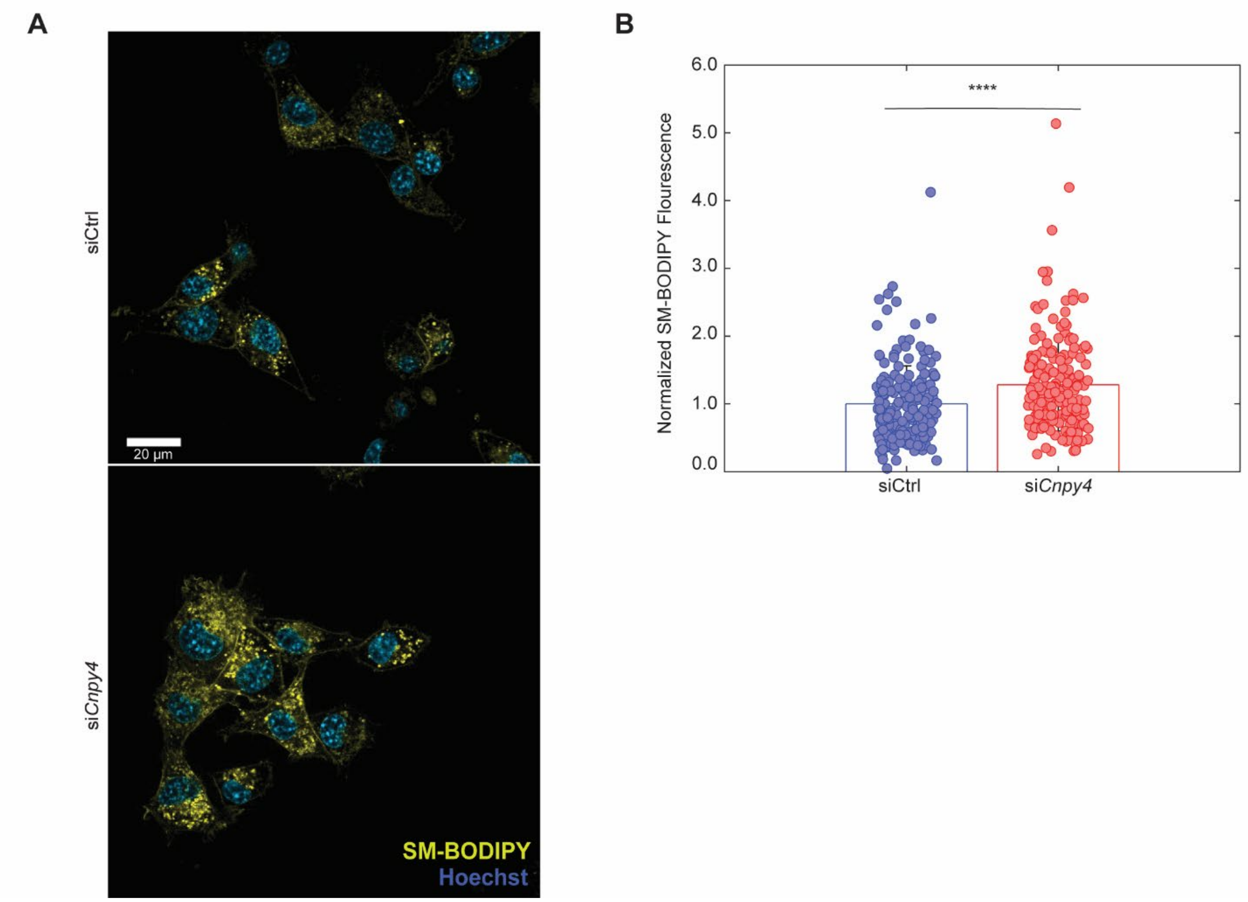


**Figure S6 Loss of CNPY4 decreases the rate of degradation of exogenous SM. (A)** Representative confocal images of live NIH3T3 cells incubated with 5 µg/mL BODIPY™ FL C12-SM for two hours prior to imaging and treated with either non-targeting siRNA control (siCtrl) or si*Cnpy4.* Hoechst 33342 was added to the medium before imaging to visualize nuclei. Scale bar corresponds to 20 µm. **(B)** Quantification of total BODIPY intensity in cells imaged as described in (A). Values were normalized to the siCtrl mean, with data combined from 4 independent experiments (n = 194 for siCtrl and n=191 for si*Cnpy4*). Statistical significance was determined using a two-sided unpaired Welch's t-test, with ****$p<0.0001$.

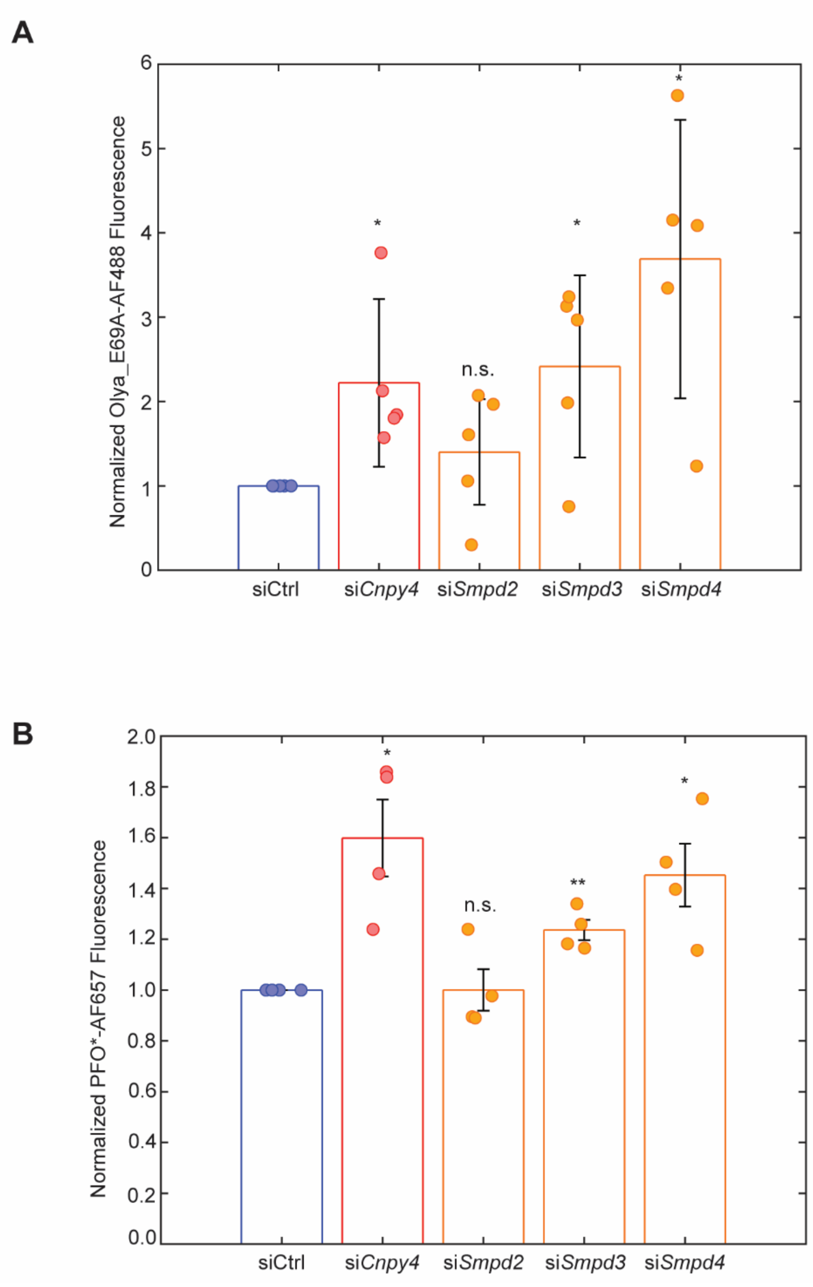


**Figure S7 Loss of nSMases increases both plasma membrane SM and free cholesterol levels. (A-B)** FACS analysis of NIHT3T3 cells treated with either non-targeting siRNA control (siCtrl), si*Cnpy4*, si*Smpd2* (nSMase1), si*Smpd3* (nSMase2), or si*Smpd4* (nSMase3), and stained with either OlyA_E69A-AF488 (total SM) in (A) or PFO*-AF647 (free cholesterol) in (B) 72 h after siRNA treatment. Data were normalized to mean value of control cells. Error bars represent mean ± SD of 4 independent trials for OlyA_E69A-AF488 and 5 independent trials for PFO*-AF647. Statistical significance was determined using a using a one-sample t-test with a null hypothesis that the mean equals 1, with *$p<0.05$, **$p<0.01$, n.s. $p>0.05$.

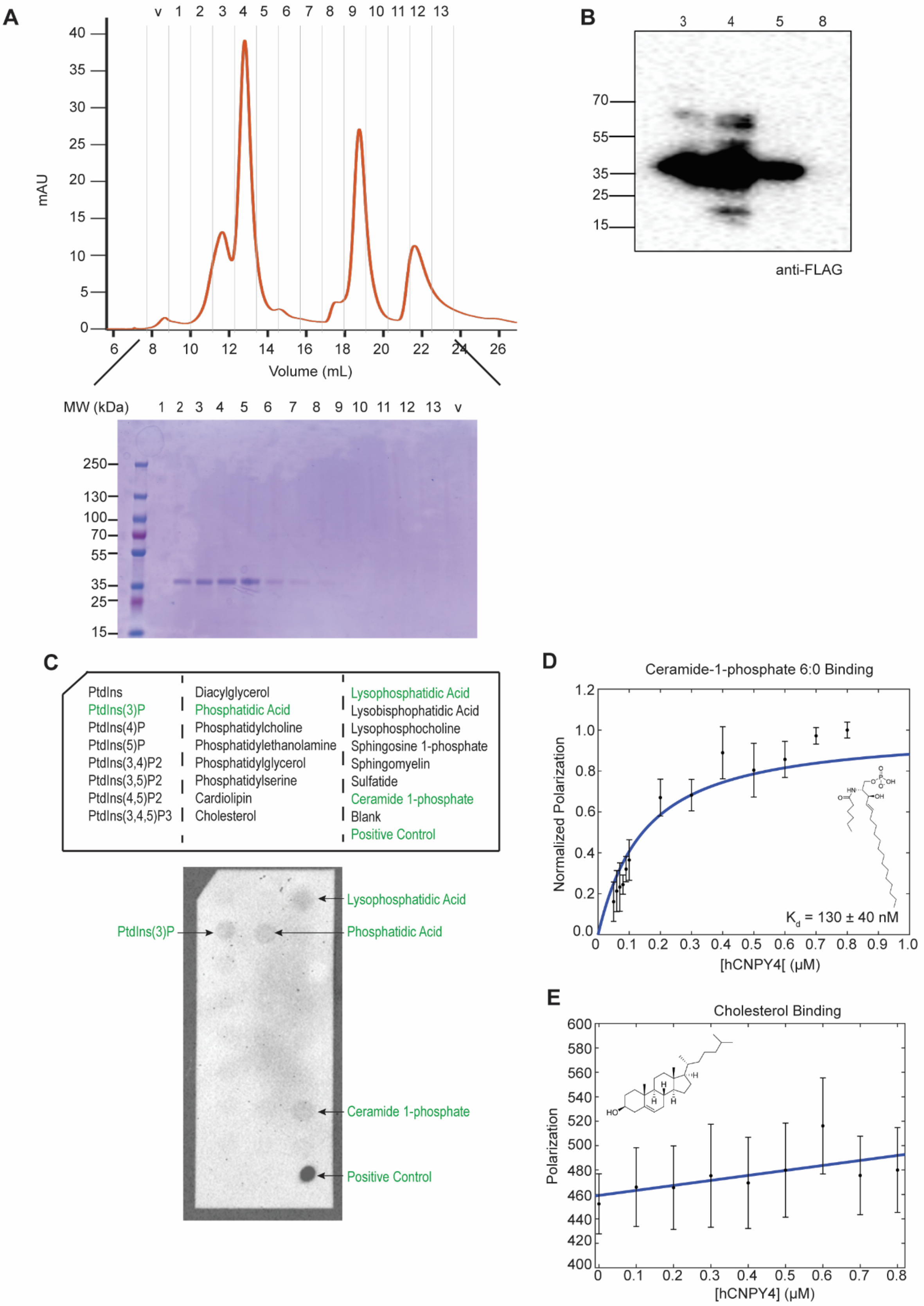
A
v 1 2 3 4 5 6 7 8 9 10 11 12 13
mAU
Volume (mL)
MW (kDa)
1 2 3 4 5 6 7 8 9 10 11 12 13 v
250
130
100
70
55
35
25
15
B
3 4 5 8
70
55
35
25
15
anti-FLAG
C
PtdIns
PtdIns(3)P
PtdIns(4)P
PtdIns(5)P
PtdIns(3,4)P2
PtdIns(3,5)P2
PtdIns(4,5)P2
PtdIns(3,4,5)P3
Diacylglycerol
Phosphatidic Acid
Phosphatidylcholine
Phosphatidylethanolamine
Phosphatidylglycerol
Phosphatidylserine
Cardiolipin
Cholesterol
Lysophosphatidic Acid
Lysobisphophatidic Acid
Lysophosphocholine
Sphingosine 1-phosphate
Sphingomyelin
Sulfatide
Ceramide 1-phosphate
Blank
Positive Control
Lysophosphatidic Acid
PtdIns(3)P
Phosphatidic Acid
Ceramide 1-phosphate
Positive Control
D
Ceramide-1-phosphate 6:0 Binding
Normalized Polarization
K$_d$ = 130 ± 40 nM
[hCNPY4[ (μM)
E
Cholesterol Binding
Polarization
[hCNPY4] (μM)

**Figure S8 Lipid binding by a recombinant hCNPY4 *in vitro*. (A)** Size exclusion chromatography (SEC) trace of the purification of the recombinant N-terminally 3xFLAG-tagged hCNPY4 and the accompanying Coomassie-stained gel. Numbers indicate the fractions sampled via Coomassie, with "v" indicating the void. **(B)** Anti-FLAG western blot of the indicated fractions from the SEC trace in (A). **(C)** hCNPY4 binding to lipids immobilized on a membrane as part of a protein-lipid blot overlay assay. Schematic representation of lipids included in the assay, with positive results in green; positive control is hCNPY4 spotted onto the membrane prior to blocking (top). Representative western blot (bottom). Membrane incubated with 8 μM of hCNPY4. **(D)** Fluorescence polarization (FP) of hCNPY4 binding to 25 nM NBD labeled 6:0 ceramide-1-phosphate (lipid shown in insert without the fluorophore). Data normalized such that the max concentration is 1 and the no protein concentration is 0. Data points represent the mean ± SEM from 3 independent experiments, with technical triplicate averaged per experiment considered one replicate. See Methods Section for details of the fit and $K_d$ determination. **(E)** FP of hCNPY4 binding to 50 nM BODIPY labeled cholesterol (lipid shown in insert without the fluorophore). Data points represent the mean ± SEM from 3 independent experiments, with technical triplicate averaged per experiment considered one replicate. Best fit line is shown, demonstrating the lack of binding.

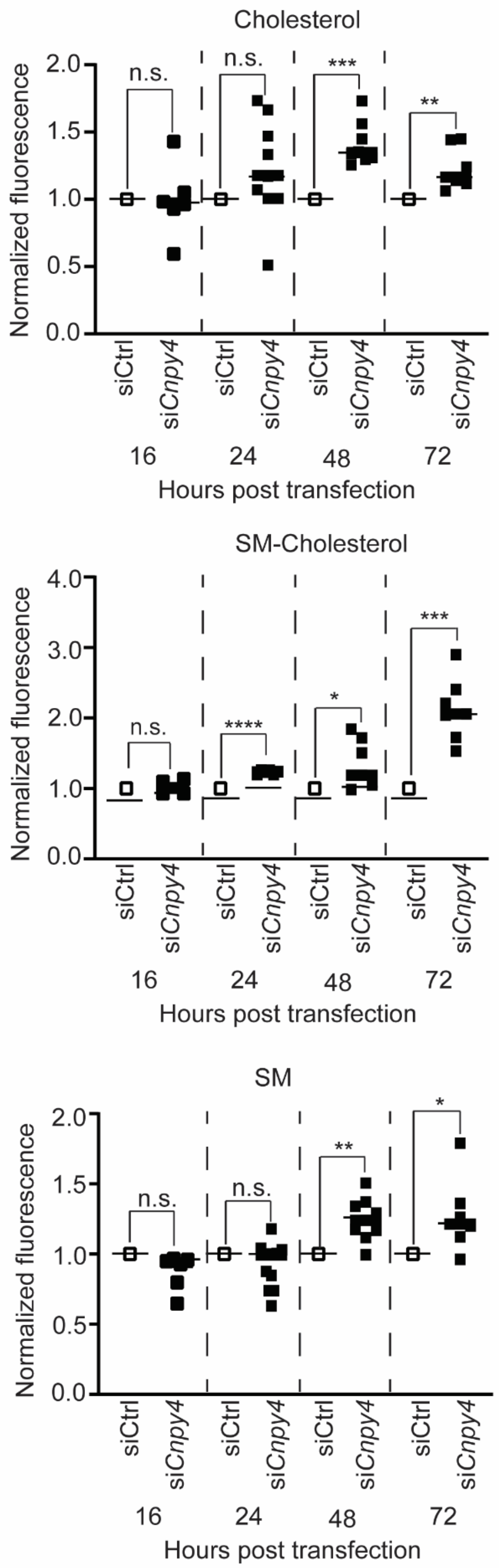
Cholesterol
Normalized fluorescence
2.0
1.5
1.0
0.5
0.0
n.s.
n.s.
***
**
siCtrl
siCnpy4
16
24
48
72
Hours post transfection
SM-Cholesterol
Normalized fluorescence
4.0
3.0
2.0
1.0
0.0
n.s.
****
*
***
siCtrl
siCnpy4
16
24
48
72
Hours post transfection
SM
Normalized fluorescence
2.0
1.5
1.0
0.5
0.0
n.s.
n.s.
**
*
siCtrl
siCnpy4
16
24
48
72
Hours post transfection

**Figure S9 Effect of *Cnpy4* knockdown on plasma membrane SM and free cholesterol levels over time.** FACS analysis of NIH3T3 cells treated with siCtrl or si*Cnpy4* and stained with PFO*-AF657 (cholesterol, top), OlyA-AF488 (SM bound to cholesterol, middle), and OlyA_E69A-AF488 (all SM, bottom) after the indicated length of siRNA treatment. Squares represent individual biological replicates normalized to the mean of the corresponding siCtrl DMSO condition, with the solid bar representing the mean of all experiments. Data were collected from 3 independent experiments for 16 h timepoint and from 5 independent experiments for 24, 48 and 72 h timepoints, with 3,000 – 10,000 live cells analyzed per one point. Statistical significance determined using a one-sample t-test with a null hypothesis that the mean equals 1, with *$p<0.05$, **$p<0.01$, ***$p<0.001$, n.s. $p>0.05$.

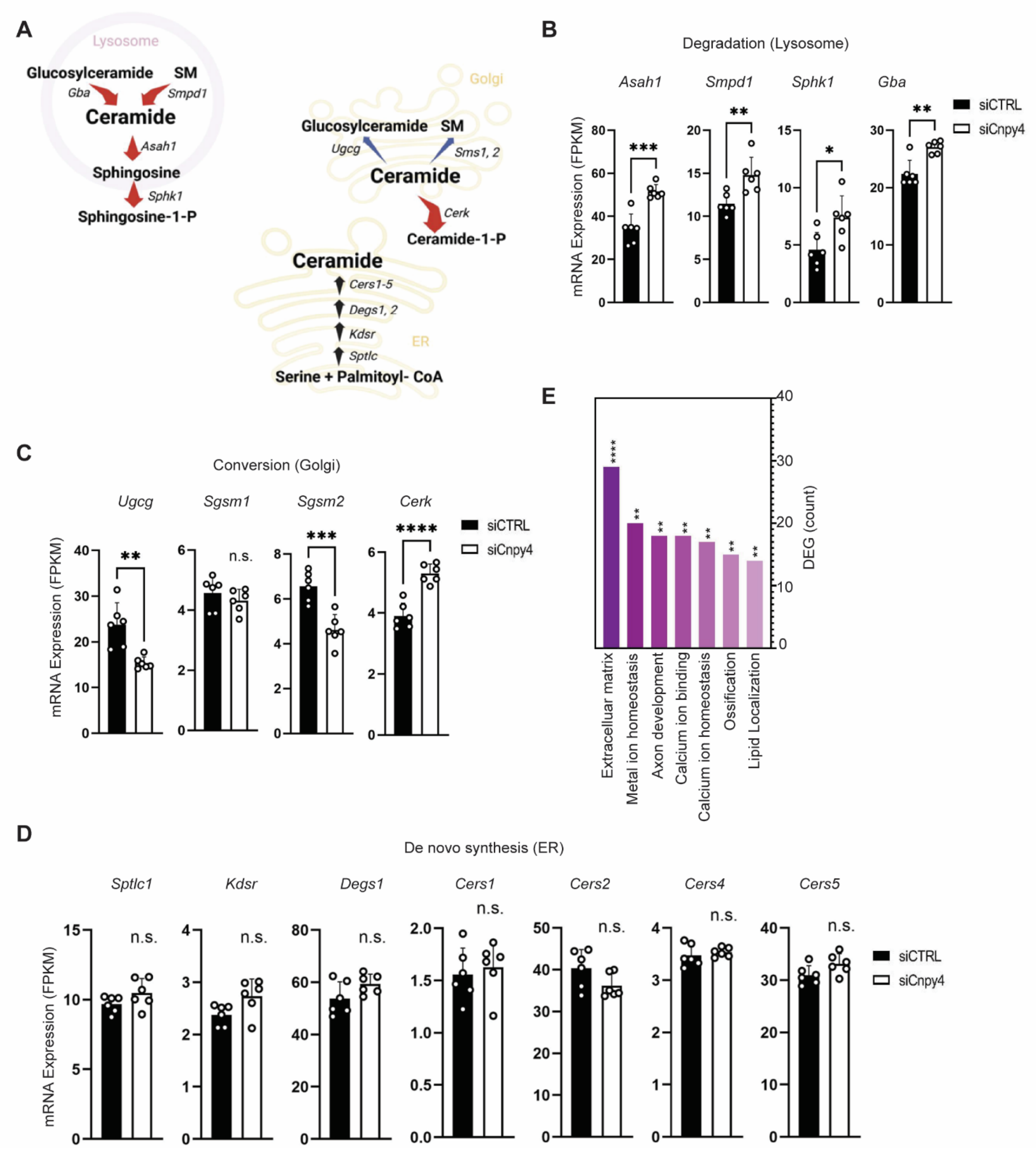


**Figure S10 Bulk RNAseq analysis of *Cnpy4* knockdown-dependent effects.** Bulk RNAseq analysis comparing NIH3T3 cells treated with either a non-targeting siRNA control (n = 6) or si*Cnpy4* (n = 6), and harvested for analysis 72 h after siRNA treatment. **(A)** Cartoon overview of pathways involved in ceramide synthesis, degradation, and conversion. Red arrows represent genes with increased expression in si*Cnpy4* treated cells, blue arrows genes with decreased expression, and black arrows with unchanged expression compared to siRNA control cells. **(B-E)**

mRNA expression of the genes highlighted in (A). Error bars represent mean ± SEM, and samples were collected from 3 independent experiments with 2 biological replicates each. (B) Genes involved in lysosomal degradation, (C) Golgi conversion to other sphingolipids, and (D) ER *de novo* synthesis. **(E)** GO analysis showing classes of genes with the largest number of differentially expressed genes (DEGs). Statistical significance was determined using DESeq2 with p-values adjustment for controlling the false discovery rate using Benjamini and Hochberg's approach, with *p<0.05, **p<0.01, ***p<0.001, ****p<0.0001, n.s. p>0.05.
